# CD36 homologs determine microbial resistance to the Lyme disease spirochete

**DOI:** 10.1101/2022.02.09.479763

**Authors:** Anya J. O’Neal, Nisha Singh, Iain S. Forrest, Agustin Rolandelli, Xiaowei Wang, Dana K. Shaw, Brianna D. Young, Sukanya Narasimhan, Shraboni Dutta, Greg A. Snyder, Liron Marnin, L. Rainer Butler, Sourabh Samaddar, M. Tays Mendes, Francy E. Cabrera Paz, Luisa M. Valencia, Eric J. Sundberg, Erol Fikrig, Utpal Pal, David J. Weber, Ron Do, Joao H.F. Pedra

## Abstract

Pattern recognition receptors sense pathogens in arthropods and mammals through distinct immune processes. Whether these molecules share a similar function and recognize the same microbe in evolutionarily distant species remain ill-defined. Here, we establish that the CD36 superfamily is required for *Borrelia burgdorferi* resistance in both the arthropod vector and humans. Using the blacklegged tick *Ixodes scapularis* and an electronic health record-linked biobank, we demonstrate that CD36 members elicit immunity to the Lyme disease spirochete. In ticks, the CD36-like protein Croquemort recognizes lipids and initiates the immune deficiency and jun N-terminal kinase pathways against *B. burgdorferi*. In humans, exome sequencing and clinical information reveal that individuals with *CD36* loss-of-function variants have increased prevalence of Lyme disease. Altogether, we discovered a conserved mechanism of anti-bacterial immunity.

**One Sentence Summary:** Lipid receptors belonging to the CD36 superfamily exhibit a shared immune function in both ticks and humans.

## Introduction

Host defense in metazoans relies on immune sensors known as pattern recognition receptors (*1*). These proteins identify molecular patterns derived from microorganisms or danger signals, leading to microbial resistance and/or tolerance (*2, 3*). This molecular archetype has been extensively explored for single host-microbe relationships. However, whether evolutionarily distant hosts share a common genetic superfamily that recognize the same microorganism cycling between species remain largely unexplored. These microbes comprise zoonotic pathogens present in animal reservoirs and livestock, which can readily crossover to humans (*4*). They also consist of pathogens that shuffle between wild animals and arthropod vectors in a sylvatic cycle, causing disease in individuals (*5*).

In North America, the blacklegged tick *Ixodes scapularis* is the primary arthropod vector for microbes that cause illness, including the Lyme disease spirochete *Borrelia burgdorferi* (*6, 7*). *B. burgdorferi* manipulates host processes for survival during infection (*6, 7*) and its lipids and lipoproteins have been shown to activate immune pathways (*8, 9*). How lipid receptors sense the presence of *B. burgdorferi* in both the arthropod vector and humans remains elusive. Here, we identified CD36 molecules as immune receptors responsible for recognizing *B. burgdorferi*. We characterized an *I. scapularis* homolog of Croquemort (Crq), a CD36-like molecule originally identified in *Drosophila*. We report that Crq is a receptor for immunostimulatory lipids and has critical roles in feeding, molting, and the tick immune response against *B. burgdorferi*. Additionally, we investigated how the human homolog CD36 contributes to immunity against *B. burgdorferi* by leveraging linked exome sequence and electronic health record data from a large-scale biobank.

## Results

### *I. scapularis* Crq is a receptor for the infection-derived lipid POPG

*Borrelia* spp. hijack and manipulate host lipids for their membranes and surface molecules (*10*). To identify lipids that may be important for tick infection, we first stimulated the *I. scapularis* cell line ISE6 with *B. burgdorferi* and performed a lipid analysis. Notably, a significant increase in the lipid 1-palmitoyl-2-oleoyl-*sn*-glycero-phosphoglycerol (POPG) was detected in tick cells after microbial stimulation (Fig. 1A and table S1). This observation was consistent with prior literature indicating that POPG is derived from bacterial infection and stimulates the immune system of ticks (*11*). Next, we used biotinylated POPG to pull down interacting proteins from tick cell lysates and eluted moieties for mass spectrometry identification. We filtered molecular hits by the gene ontology (GO) term “membrane” and the keyword “receptor” to identify potential transmembrane lipid receptors (table S2). One hit was annotated as “scavenger receptor class B type I (SR-B1)”. A basic local alignment search tool (BLAST) analysis revealed that this molecule is homologous to the *Drosophila* receptor Crq, a member of the CD36 superfamily (*12, 13*). The CD36 superfamily is an ancient group of lipid scavenger receptors with roles in metabolism and immunity (*14-16*). In *Drosophila*, Crq is a receptor of apoptotic cells and is involved in lipid uptake, antibacterial immunity and jun N-terminal kinase (JNK) activation (*13, 17, 18*). Lipid scavenger receptors have also been shown to contribute to *B. burgdorferi* phagocytosis in mammals (*19*). Thus, we focused on this protein, which we named *I. scapularis* Crq (Fig. 1B) (LOC8027712).

**Figure 1.**
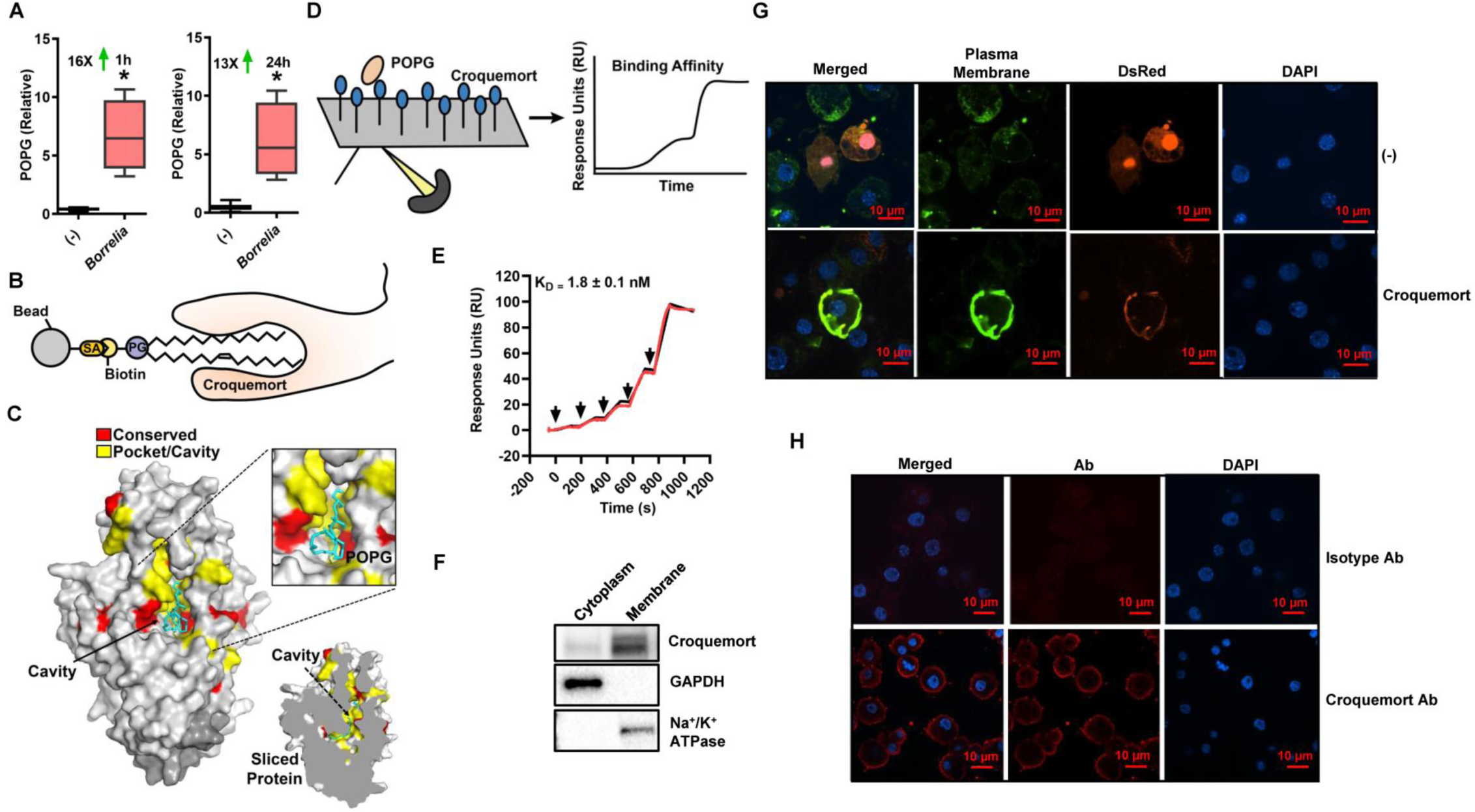
*I. scapularis* Crq is a receptor for the infection-derived lipid POPG. **A**. POPG levels in ISE6 cells stimulated with *B. burgdorferi* (MOI 500) for 1 hour and 24 hours relative to unstimulated (-) cells. Lipids were measured by mass spectrometry. Results are represented as box-and-whisker plots (*n*=4-6). **B**. Diagram of pulldown approach. **C**. Modeling of the *I. scapularis* Crq ectodomain using Phyre2 (*24*). The model exhibits a canonical scavenger receptor type fold with a large open cavity. Conserved residues are labeled in red and potential pocket residues are labeled in yellow. The POPG and the Crq homology model placed POPG (cyan) within the interior of this predicted cavity in 8 out of 9 Autodock trials (*26*). **D-E**. Representative surface plasmon resonance sensorgram (red) from immobilized recombinant Crq ectodomain (Crq-His) binding to POPG injected at 12.5 nM, 25 nM, 50 nM, 100 nM, and 200 nM. The data was fit to a single cycle kinetic model (black), and the dissociation constant (K_D_) was determined by k_off_/k_on_ to be 1.8 +/- 0.1 nM from duplicate experiments. Arrows denote time of lipid injection. **F**. Subcellular fractionation of ISE6 cells. **G**. Ectopic expression of DsRed-tagged Crq in the IDE12 tick cell line. IDE12 cells were nucleofected with plasmid containing Crq-DsRed or empty vector (-). Green - plasma membrane; blue – DAPI (4’,6-diamidino-2-phenylindole). Data represent one of two independent experiments. **H**. IDE12 cells were stained with the anti-Crq or isotype control antibodies. Data represent one of two independent experiments. Statistical significance was evaluated by an unpaired *t* test with Welch’s correction. *, *p*<0.05. SA=streptavidin, PG=phosphatidylglycerol, Ab=antibody; Crq=Croquemort.

CD36 molecules have two transmembrane regions, an extracellular portion with a characteristic cavity opening and two intracellular tails (*16, 20, 21*). Consistent with this, Crq possessed an ectodomain and carried two transmembrane domains, as shown through transmembrane topology prediction (*22*). Because electrostatic interactions in CD36-like molecules contribute to the association of these receptors with polyanionic ligands (*20, 23*), we next investigated the residues that favor contact between Crq and POPG. We aligned Crq with the human molecules: lysosome membrane protein 2 (LIMP-2) (Q14108) and CD36 (P16671) (fig. S1). Furthermore, we modeled the ectodomain of Crq to CD36 and performed homology comparisons between Crq and the crystal structures of LIMP-2 (PDB:4F7B) and CD36 (PDB:5LGD) using Protein Homology/Analogy Recognition Engine (Phyre) 2 (*24*) and AlphaFold Protein Structural Database (*25*) (Fig. 1C and fig. S2). Finally, we determined potential ligand-protein interactions between Crq and POPG using AutoDock (*26*). Altogether, we determined that Crq carried a canonical scavenger receptor type fold with a large open cavity or pocket where POPG was predicted to interact (Fig. 1C).

To validate our model empirically, we first expressed and purified a His-tagged Crq ectodomain (Crq-His) (fig. S3A and S3B). We then confirmed that this protein was folded using 2D nuclear magnetic resonance spectroscopy, as determined by significant dispersion of Crq ^1^H and ^15^N chemical shift values (fig. S3C). Lastly, we observed that Crq-His bound POPG in the low nanomolar range using surface plasmon resonance with single cycle kinetics (Figs. 1D and 1E). Taken together, we determined that the ectodomain of Crq bound POPG at the interface.

Mammalian CD36 and *Drosophila* Crq are plasma membrane-bound receptors (*12, 14, 15*). Next, we performed subcellular fractionation to confirm the membrane localization of Crq (Fig. 1F). We also developed a protocol for ectopic expression to visualize Crq localization within the cell (fig S4A and S4B). Prior to this work, expression of genes in tick cells was not technically feasible because they carry fastidious requirements for genetic manipulation (*27*). We observed that Crq localized with the plasma membrane in the hemocyte-like IDE12 and the neuronal-like ISE6 *I. scapularis* cell lines (Fig. 1G and fig. S4C). Crq was also detected on the plasma membrane of IDE12 cells using a Crq-specific antibody (Fig. 1H). Thus, these data indicate that Crq is a plasma membrane receptor for infection-derived lipids.

### *I. scapularis* Crq restricts *B. burgdorferi* colonization through the IMD and JNK pathways

The central dogma of arthropod immunity states that Gram-negative bacteria activate the immune deficiency (IMD) pathway, a nuclear factor (NF)-κB signaling network, through the cell wall component diaminopimelic peptidoglycan (*28, 29*). Notably, the IMD signaling relay occurs distinctively in *Drosophila* when compared to other non-insect arthropods. For instance, ticks possess core intracellular components of the “canonical” IMD pathway, such as the NF-κB transcription factor Relish, which restricts colonization of *B. burgdorferi* (*11, 30*). However, genome and functional analyses revealed that ticks lack key upstream components, including transmembrane peptidoglycan recognition proteins and the adapter molecules IMD and FADD (*11, 30-33*). Cell signaling in *I. scapularis* is relayed through “non-canonical” molecules, including p47 (*30*). Importantly, the tick IMD pathway is activated by lipid molecular patterns, such as POPG (*11*).

Given the ability of CD36 to bind immunogenic lipids, we speculated that Crq may be the lipid receptor for the *I. scapularis* IMD pathway. To evaluate this hypothesis, *I. scapularis* IDE12 cells were transfected with small interfering RNA (siRNA) to silence *crq* expression (Fig. 2A). RNA interference (RNAi) remains the gold standard for disruption of gene function in tick cells, as clustered regularly interspaced short palindromic repeats (CRISPR) technology has not yet been established in this system (*27*). Next, we stimulated *I. scapularis* IDE12 cells with the lipid POPG. POPG stimulation led to cleavage of the IMD-specific NF-κB molecule Relish and phosphorylation of JNK (Fig. 2B). However, *crq* silencing reduced Relish processing and altered JNK phosphorylation (Figs. 2B and 2C). Following the initiation of IMD signaling in *Drosophila*, various proteins are recruited to the molecular scaffold, including transforming growth factor-β activated kinase 1 (TAK1). TAK1 promotes JNK activation in parallel to NF-κB signaling (*34, 35*). To ascertain whether *I. scapularis* TAK1 was required for POPG-mediated immune activation, we transfected IDE12 cells with *tak1* siRNA (Fig. 2D). Importantly, TAK1 was required for both the accumulation of N-Rel and JNK phosphorylation during POPG stimulation (Figs. 2E and 2F).

**Figure 2.**
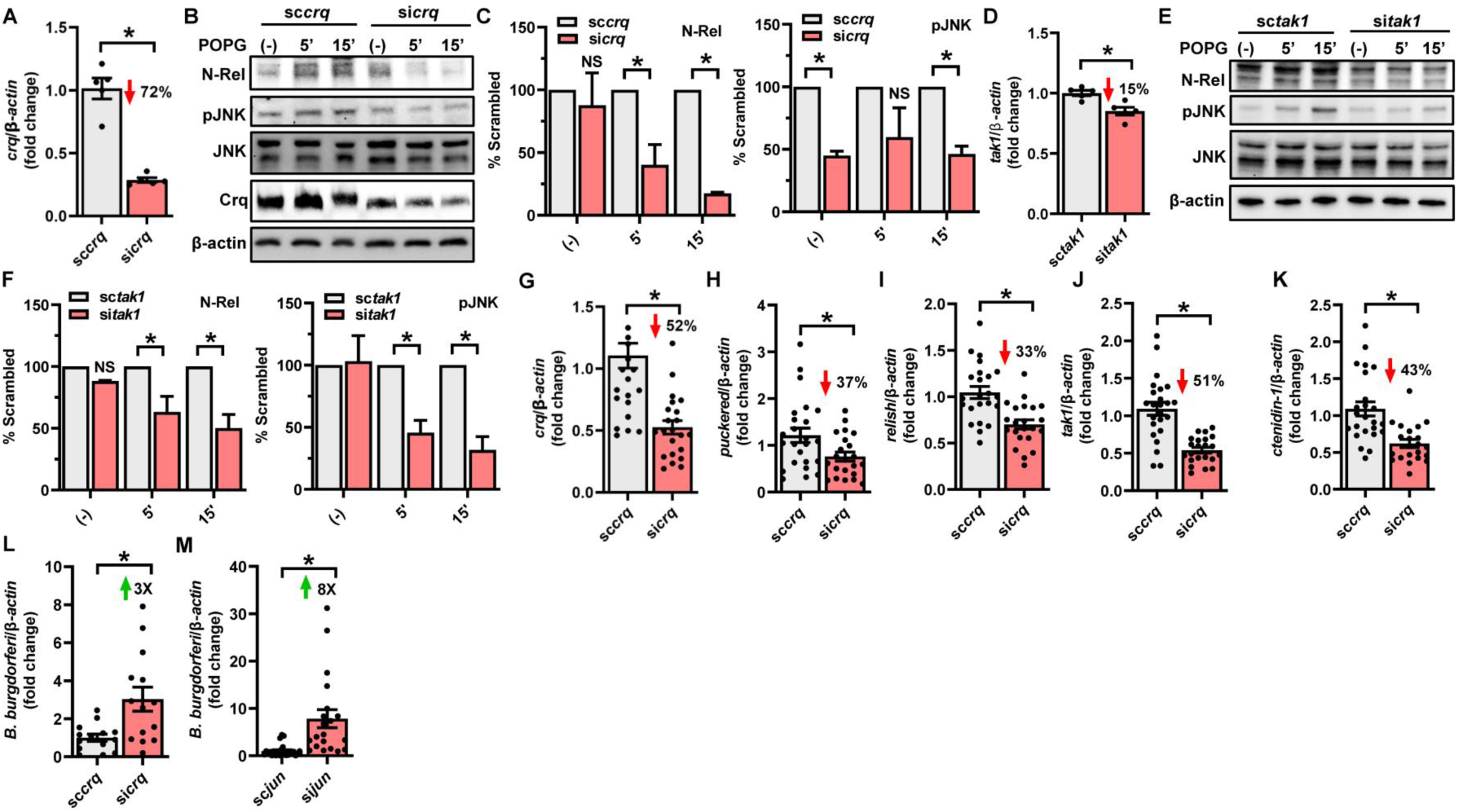
*I. scapularis* Crq restricts *B. burgdorferi* acquisition through activation of the IMD and JNK pathways. **A**. *crq* silencing efficiency in IDE12 cells. Cells were transfected with *crq* siRNA (si*crq*) or scrambled RNA (sc*crq*) (*n*=5). **B and C**. *crq* silenced or scrambled control IDE12 cells were stimulated with 10 ng/ml POPG for the indicated time points. Normalized data for si*crq* were divided by the corresponding control values and expressed as a percentage of sc*crq*. N-rel values are normalized to actin and pJNK values are normalized to JNK (p52). Western blot bands were quantified using ImageJ. Western blot images show one representative experiment (*n*=2-3). **D**. *tak1* silencing efficiency in IDE12 cells. Cells were transfected with *tak1* siRNA (si*tak1*) or scrambled RNA (sc*tak1*) (*n*=5). **E and F**. *tak1* silenced or scrambled control IDE12 cells were stimulated with 10 ng/ml POPG for indicated time points. Normalized data for si*tak1* were divided by the corresponding control values and expressed as a percentage of sc*tak1*. N-rel values are normalized to actin and pJNK values are normalized to JNK (p52). Western blot bands were quantified using ImageJ. Western blot images show one representative experiment (*n*=2). **G**. *crq* silencing efficiency in ticks microinjected with si*crq* and sc*crq*. Ticks were allowed to feed to repletion (up to five days) (*n*=22-24). **H-K**. Expression of JNK- and IMD-related immune genes in fully repleted si*crq* and sc*crq* ticks (*n*=21-24). **L**. Bacterial acquisition in si*crq* and sc*crq* ticks fed on mice infected with *B. burgdorferi. B. burgdorferi* burden was quantified by *recA* expression (*n*=14). **M**. Bacterial acquisition in si*jun* and sc*jun* ticks fed on mice infected with *B. burgdorferi. B. burgdorferi* burden was quantified by *recA* expression (*n*=20-24). Results are represented as means +/- SE. At least two biological replicates were performed. Statistical significance was evaluated by an unpaired *t* test with Welch’s correction (A, D, G-M) or a two-way ANOVA post-hoc Sidak test for multiple comparisons (C, F). *, *p*<0.05. (-)=unstimulated, N=Rel=cleaved Relish, pJNK=phosphorylated JNK.

JNK signaling activates the transcription factor Jun and the phosphatase *puckered* can be used as a genetic readout for JNK activation (*36*). The *I. scapularis* genome encodes a homolog of *puckered* (annotated as “dual specificity protein phosphatase 10”) which was downregulated during *jun* silencing in nymphal ticks (fig. S5). To determine whether Crq regulated *puckered*, non-engorged *I. scapularis* nymphs were microinjected with siRNA to silence *crq* expression (Fig. 2G). Injected *I. scapularis* nymphs were allowed to attach and feed until repletion. Notably, *crq* silencing not only decreased the expression of *puckered* (Fig. 2H), but also the IMD signaling components *relish* (Fig. 2I) and *tak1* (Fig. 2J).

Activation of the *Drosophila* IMD pathway results in the transcription of specific antimicrobial peptides (AMPs) (*34, 35*). Unlike *Drosophila*, arachnids have been shown to constitutively express and store their AMPs within hemocyte granules (*37, 38*). Ctenidins are a family of glycine-rich AMPs identified from the spider *Cupiennius salei* that exhibit activity against Gram-negative bacteria (*37, 38*). *I. scapularis* carried several peptides annotated as “ctenidin-1-like” (fig. S6). Hence, we investigated whether an *I. scapularis* ctenidin-1 was specifically regulated by the tick IMD pathway. Electrophoretic mobility shift assay determined that Relish binds to the *ctenidin-1* promoter (figs. S7A and S7B). Furthermore, transcriptional silencing of *relish* resulted in decreased expression of *ctenidin-1* both *in vitro* and *in vivo* (figs. S7C and S7D). Conversely, silencing of the Toll-specific NF-κB molecule, *dorsal*, did not affect *ctenidin-1* expression (figs. S7E and S7F). Importantly, *crq* silencing reduced the expression of *ctenidin-1* (Fig. 2K), suggesting specific regulation of *ctenidin-1* by the *I. scapularis* IMD network. We then investigated whether this *I. scapularis* immune signaling relay impacted *B. burgdorferi* infection in ticks. Indeed, *crq* silencing in nymphs resulted in significantly higher *B. burgdorferi* burden (Fig. 2L and fig. S8). Likewise, *jun*-silenced ticks (fig. S5) acquired significantly more *B. burgdorferi* compared to tick-treated scrambled siRNA controls (Fig. 2M). Altogether, our findings suggested that Crq led a concerted action by the IMD and the JNK signaling networks to promote resistance against the spirochete *B. burgdorferi* in ticks (fig. S9).

### *I. scapularis* Crq regulates tick feeding and molting

CD36 molecules have various functions, including microbial sensing, lipid scavenging and cell adhesion (*14, 15*). Notably, immunity and fitness are physiologically connected in arthropods (*39*) and hematophagous vectors depend on lipid metabolism for development (*40*). Thus, we allowed nymphs to feed until repletion and measured *crq* expression over time. We found that *crq* was upregulated during early tick feeding but not during full engorgement (fig. S10A). To determine if Crq contributed to tick fitness, we silenced *crq* in nymphs (Fig. 2G) and allowed them to feed until repletion. We did not observe a difference in attachment between treatments (fig. S10B). However, the average weight of *crq* silenced ticks was significantly lower than scrambled controls (Fig. 3A), indicating that Crq has an important role in arthropod feeding.

**Figure 3.**
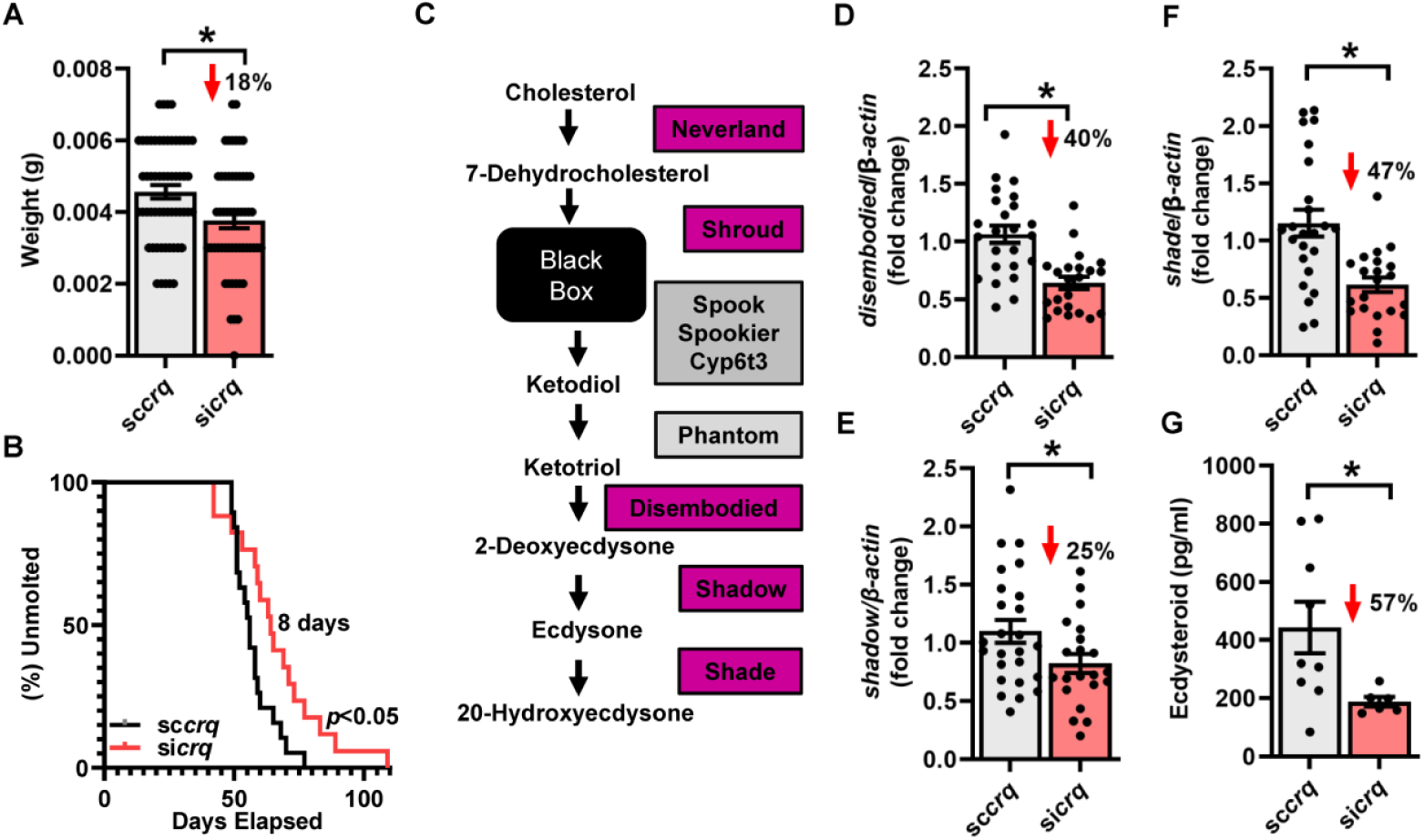
*I. scapularis* Crq regulates tick feeding and molting. **A**. Weight of fully repleted nymphs microinjected with *crq* siRNA (si*crq*) or scrambled RNA (sc*crq*) (*n*=55-58). **B**. Molting curve of si*crq* and sc*crq* nymphs that successfully molted into adults. Nymphs were allowed to feed until repletion and time of molting was monitored (*n*=17-19). **C**. Diagram of 20-hydroxyecdysone synthesis in *D. melanogaster*. Enzymes shaded in magenta were found in the *I. scapularis* genome, enzymes shaded in gray were not found. **D-F**. Expression of ecdysteroid synthesis enzymes (“Halloween genes”) in fully repleted si*crq* and sc*crq* nymphs (*n*=21-25). **G**. Concentration of ecdysteroids in fully repleted si*crq* and sc*crq* nymphs (*n*=6-9). Results are represented as means +/- SE. At least two biological replicates were performed. Statistical significance was evaluated by an unpaired *t* test with Welch’s correction (A, D-G) or a Log-rank (Mantel-Cox) test (B). *, *p*<0.05. g=grams, pg=picograms, ml=milliliters.

Ticks develop into adults through an incomplete metamorphosis, a process that is accomplished by molting or shedding of the exoskeleton (*41, 42*). Arthropod molting is regulated by ecdysteroids, such as 20-hydroxyecdysone (20E) and ponasterone A, which are synthesized from dietary cholesterol (*41, 42*). Because CD36 facilitates the transport of lipids, including cholesterol into mammalian cells, we posited that *crq* silencing would disrupt the molting process in *I. scapularis*. Thus, we allowed fully repleted *crq* silenced nymphs to molt into adults and measured their molting capacity. Metamorphosis to adults was comparable between silenced and control ticks (fig. S10C). However, molting in *crq* silenced ticks was delayed by 8 days when compared to scrambled controls (Fig. 3B). We then hypothesized that the delay of molting in silenced ticks was due to defective ecdysteroid production. In *Drosophila*, cholesterol is converted into 20E by a family of enzymes encoded by the “Halloween” genes (*42, 43*) (Fig. 3C). *I. scapularis* carries homologs of most ecdysteroid enzymes, excluding the enzyme encoded by *phantom*, which converts 5β-ketodiol to 5β-ketotriol via hydroxylation of carbon 25 on the tetracyclic steroidal backbone (*44, 45*) (Fig. 3C). We then quantified the expression of ecdysteroid-related genes in *crq* silenced nymphs and observed decreased transcription of *disembodied* (Fig. 3D), *shadow* (Fig. 3E) and *shade* (Fig. 3F), but not the ecdysone receptors *ecr* and *usp* (ultraspiracle) (figs. S10D and S10E). Supporting these findings, *crq* silenced nymphs produced fewer ecdysteroids compared to controls (Fig. 3G). Taken together, these data suggest that Crq maintains tick fitness through feeding and regulation of molting enzymes.

### Loss-of-function variants in the *CD36* gene associate with Lyme disease diagnosis in a population-based biobank

Because the CD36 superfamily is evolutionarily conserved, we reasoned that immune recognition of *B. burgdorferi* may share a common protein sensor in both ticks and humans. To investigate this, we ascertained whether the CD36 gene family is associated with human Lyme disease using the electronic health record-linked biobank Bio*Me* (Fig. 4A) (*46*). *CD36* was less genetically constrained compared to the other family members *SCARB1* (SR-BI) or *SCARB2* (LIMP-2) according to the Genome Aggregation Database (gnomAD) (table S3) (*47*). Therefore, we identified individuals with rare loss-of-function (LoF) variants in *CD36*. Out of 28,877 individuals with exome sequence and electronic health record data, 394 (1.4%) possessed at least one rare LoF variant allele (Fig. 4A and table S4). In total, three *CD36* LoF variants were identified using the Variant Effect Predictor (Fig. 4B and table S4).

**Figure 4.**
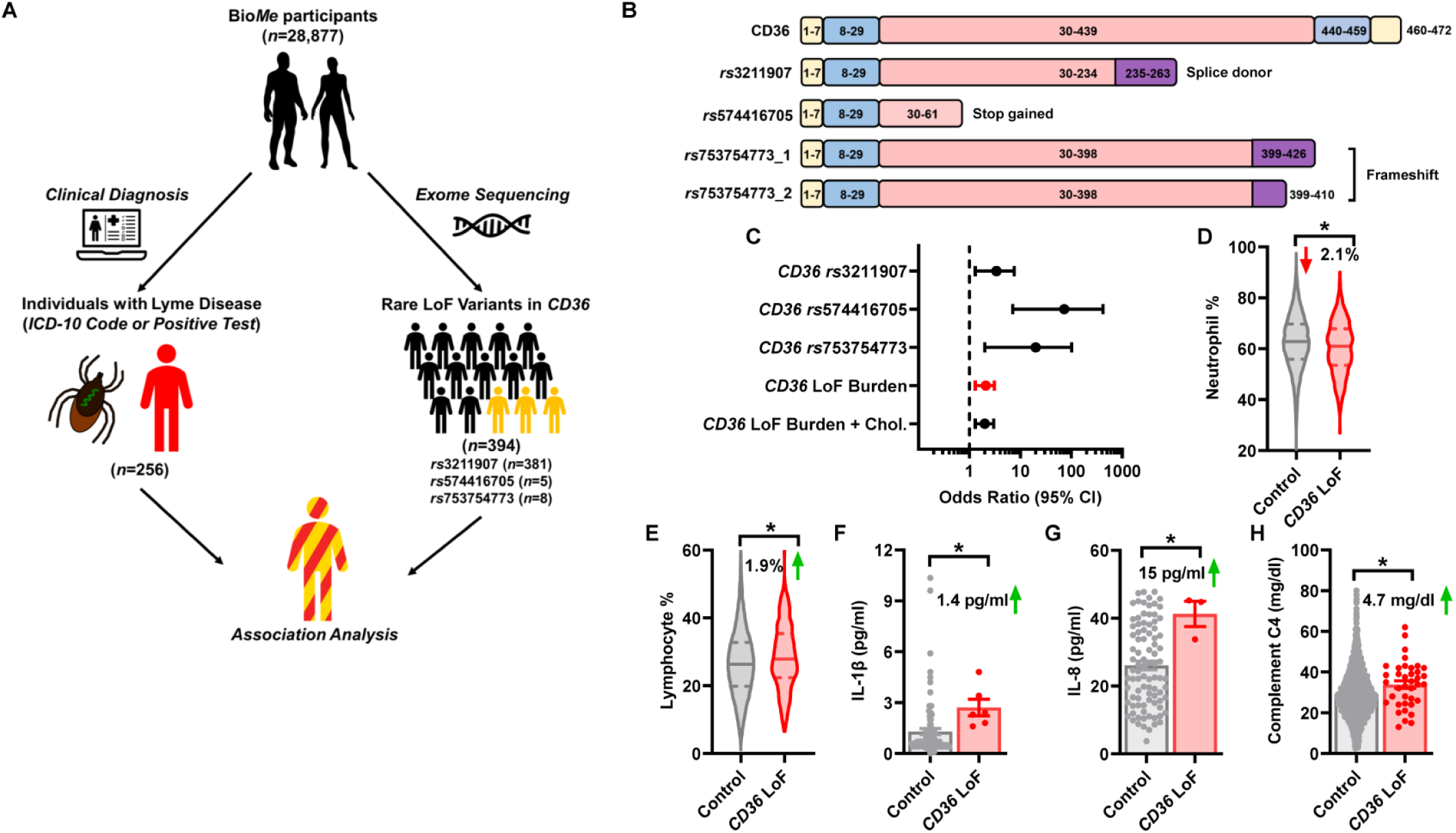
Evaluation of *CD36* loss-of-function burden association with Lyme disease in 28,877 participants with exome and electronic health records. **A**. A cohort of 28,877 individuals from Bio*Me*, a health system-based biobank in New York City, had linked exome sequence and electronic health record (EHR) data. Three rare loss-of-function (LoF) variants in *CD36* were identified in 394 individuals with annotations of frameshift, splice acceptor/donor, stop gained/lost, or start lost using Variant Effect Predictor (VEP). Allele counts for these three variants were aggregated into a single *CD36* LoF burden score for individuals. Cases of Lyme disease were identified in 256 individuals by the presence of a physician-documented International Classification of Diseases-Clinical Modification 10 (ICD-10) diagnosis code of A69.2 or a positive *B. burgdorferi* antibody or PCR laboratory test. The association of *CD36* LoF burden with Lyme disease was evaluated using multivariable regression with Firth’s penalized likelihood, adjusted for age, sex, body mass index, and 10 genetic principal components. **B**. Comparison of predicted *CD36* LoF variants at the protein level. Altered amino acid sequences due to frameshift or splice donor mutations are labeled in purple. *rs*753754773 consists of two possible protein outcomes due to an insertion or deletion at the same position. Yellow=intracellular tails, blue=transmembrane domain, pink=ectodomain. **C**. Forest plot on a base-10 logarithmic scale depicts the adjusted odds ratio and 95% confidence interval (CI) of Lyme disease associated with each of the three *CD36* LoF variants, the *CD36* LoF burden score (red), and the *CD36* LoF burden score additionally adjusted for total cholesterol level. **D**. Violin plots of the distribution of neutrophil percentage in individuals without a *CD36* LoF variant (control; *n*=23,947) versus individuals with a *CD36* LoF variant (*CD36* LoF; *n*=349). **E**. Violin plots of the distribution of lymphocyte percentage in individuals without a *CD36* LoF variant (control; *n*=23,984) versus individuals with a *CD36* LoF variant (*CD36* LoF; *n*=351). **F**. IL-1β levels in individuals without a *CD36* LoF variant (control; *n*=87) versus individuals with a *CD36* LoF variant (*CD36* LoF; *n*=6). **G**. IL-8 levels in individuals without a *CD36* LoF variant (control; *n*=90) versus individuals with a *CD36* LoF variant (*CD36* LoF; *n*=3). **H**. Complement C4 levels in individuals without a *CD36* LoF variant (control; *n*=2073) versus individuals with a *CD36* LoF variant (*CD36* LoF; *n*=38). Results are represented as violin plots (D and E) or means +/- SE (F-H). Statistical significance was evaluated by an unpaired *t* test with Welch’s correction. *, *p*<0.05. Cholesterol and other laboratory levels were obtained as the median value. Thus, if an individual had their cholesterol or a laboratory test measured several times a day in the EHR, we displayed the median value. Other clinical variables were taken at baseline when the participants were first enrolled at Bio*Me*.

*CD36* LoF variants have been associated with increased cholesterol levels (*48*). To validate the functional relevance of three variants of interest, we obtained cholesterol measurements from electronic health records and calculated a *CD36* gene burden score, which aggregates allele counts across all three predicted LoF variants. We observed that *CD36* LoF burden was associated with increased total cholesterol (4.4 mg/dL increase per LoF allele; standard error =2.1 mg/dL; *p*=0.04), adjusted for statin usage and clinical covariates (age, sex, body mass index, and 10 genetic principal components) (table S5). Then, we correlated *CD36* LoF variants with Lyme disease in the study population. We identified 256 cases of Lyme disease that were physician diagnosed and/or laboratory confirmed in accordance with the Centers for Disease Control and Prevention guidelines. We performed a multivariable regression analysis of Lyme disease as a function of each *CD36* LoF variant. We found that all three *CD36* LoF variants were individually associated with Lyme disease (*p*<0.05; Fig. 4C and table S4).

We then assessed the association of *CD36* LoF burden with Lyme disease diagnosis, adjusting for clinical covariates. Notably, we found that *CD36* LoF burden was significantly associated with Lyme disease diagnosis with a 2.1-fold increased odds of diagnosis per LoF allele (95% confidence interval [CI] = 1.3-3.1; *p*= 3.1 × 10^−4^) (Fig. 4C and table S6). Hypercholesterolemia has been shown to increase pathogenicity of *B. burgdorferi* infection (*49*). To determine if genetic susceptibility to Lyme disease in individuals with *CD36* LoF variants was attributed to elevated cholesterol, we included total cholesterol levels as a covariate in our multivariable regression analysis. *CD36* LoF burden remained significantly associated with Lyme disease diagnosis with a 2.0-fold increased odds of diagnosis per increase in LoF allele (95% CI = 1.3-3.0; *p*=1.1 × 10^−3^) (Fig. 4C and table S7). This finding indicated that rare *CD36* LoF variants associate with Lyme disease for reasons other than hypercholesterolemia.

In symptomatic patients, Lyme disease typically presents with inflammatory symptoms (*6, 7*). Based on the findings described above, we ascertained whether individuals with *CD36* LoF variants had an altered immune response. Among a subset of participants with laboratory results, we found that individuals with *CD36* LoF variants had a 2.1 percentage point lower mean of neutrophils (*p*=1 × 10^−4^) (Fig. 4D) and 1.9 percentage point elevated mean of lymphocytes compared to individuals without *CD36* LoF variants (*p*=2 × 10^−4^) (Fig. 4E). Consistent with this, the effect size on leukocyte composition with *CD36* LoF was similar or greater than that observed in previous LoF studies of other genes (*51*). Individuals with *CD36* LoF variants also had higher mean levels of inflammatory markers, including the pyrogenic cytokine interleukin (IL)-1β (*p*=0.032) (Fig. 4F), the neutrophil chemotactic factor IL-8 (*p*=0.043) (Fig. 4G) and complement C4 (*p*=0.014) (Fig. 4H).

Functional studies with the human variants are not technically feasible based on their likely biogenesis, topology and biophysical features (*52*). Hence, we characterized the inflammatory potential of CD36 in mice. We isolated bone marrow-derived macrophages (BMDMs) from *Cd36*^-/-^ and wild-type (WT) C57BL/6J mice and stimulated these cells with *B. burgdorferi*. Production of proinflammatory cytokines was affected, including an increase in IL-1β, as seen in humans, as well as decreased IL-6 and keratinocyte-derived chemokine (KC) (fig. S11A). Additionally, JNK and NF-κB activation were reduced in *Cd36*^-/-^ BMDMs after 15 minutes of *B. burgdorferi* stimulation (figs. S11B-D), which is consistent with previous studies implicating CD36 in these pathways (*53, 54*). Altogether, our results reveal a significant role for *CD36* in the immune response to *B. burgdorferi*.

## Discussion

Evolution promotes a duality of uniqueness and universality (*55*). While organismal diversity is a primary outcome of natural selection within distinct environments, there is a paradoxical preservation of shared, fundamental functions across species. The universality of life is evident in ancient gene families whose features have been maintained in distant organisms. Our study shows that CD36 molecules are important for immunity to *B. burgdorferi* in species separated by over 700 million years of evolutionary history. In ticks, Crq binds infection-derived lipids and relays antimicrobial signals through the IMD and JNK pathways. Receptors for these pathways were previously undefined in chelicerates due to a lack of conservation with *Drosophila* immune components. Ticks are non-model organisms and members of the subphylum Chelicerata, an ancient group of arthropods that predate dipteran insects (*56*). Thus, these data establish a primordial mechanism of immune recognition that may have broader implications for arthropod immunology. Interestingly, other tick-borne bacteria, including *Anaplasma* spp., activate the tick IMD pathway through POPG (*11*). Whether CD36 molecules are involved in the immune response to other tick microbes remains to be seen.

Biobank data revealed that individuals with rare *CD36* LoF variants have higher prevalence of Lyme disease regardless of the genetic diversity and variability among a large population in Bio*Me* (*46*). Our findings were achieved despite the multifactorial and polygenic nature of immune traits (*57*) and distinctive non-heritable factors, such as physical activity, prior infections, diet, vaccination status and psychosocial stress (*58*). Although striking, our results have some limitations. First, the epidemiological data available in Bio*Me* do not distinguish between different stages of *B. burgdorferi* infection, such as early or late disease, or post-treatment Lyme syndrome (*6, 59*). Second, electronic health records were obtained from consenting individuals in the Mount Sinai Health System, which is based in the New York City metropolitan area (table S8). While Lyme disease is highly prevalent in the northeastern United States, it is unclear whether our results are generalizable to other regions of the United States and/or endemic countries elsewhere (*6, 59*). Third, clinical and laboratory data were only available in electronic health records for a subset of individuals. Future endeavors incorporating sequencing and clinical phenotyping should provide increased knowledge about the genetic epidemiology of Lyme disease. Finally, two *CD36* LoF variants had a small number of individuals, which reduced the power for discovery.

In summary, our results highlighted the importance of studying the interactions between ticks and *B. burgdorferi* to uncover new insights related to Lyme disease. Recently, advances in personalized medicine enabled innovative opportunities to combat various infectious and non-infectious illnesses. Characterizing the vector-pathogen interface may aid in our understanding of human immunity in arthropod-borne illnesses. Future investigations should provide a useful blueprint for studying the epidemiology of Lyme disease.

## Acknowledgements

The authors acknowledge members of the Pedra laboratory for providing insightful discussions and colleagues for manuscript feedback. We thank Jon Skare (Texas A&M University Health Science Center) for providing the *B. burgdorferi* B31 strain, clone MSK5; Ulrike G. Munderloh (University of Minnesota) for supplying ISE6 and IDE12 tick cells; Yinghua Zhang (University of Maryland School of Medicine) for surface plasmon resonance analysis; Joseph Mauban (University of Maryland School of Medicine) for aiding in the microscopy analysis; Jonathan Oliver (University of Minnesota) for making available *I. scapularis* ticks; and Holly Hammond (University of Maryland School of Medicine) for providing administrative support.

## Author contributions

AJO conceived, designed, and carried out experiments, analyzed data, prepared figures, and wrote the paper. ISF designed and carried out experiments, analyzed data, prepared figures, and wrote the paper. NS and AR carried out experiments, analyzed data, and wrote methods. XW and DKS designed experiments, carried out experiments, and analyzed data. BDY carried out experiments, analyzed data, prepared figures, and wrote methods. SN, SD, LM, LRB, SS, MTM, FECP, and LMV carried out experiments. GAS performed protein modeling and prepared figures. DJW, EJS, UP, EF and RD supervised experiments, provided reagents, and analyzed data. JHFP conceived and designed experiments, supervised the study, prepared figures, and wrote the paper.

## Data and materials availability

All data are available in the manuscript or in the supplementary materials.

## Supplementary Materials

Materials and Methods

Figs. S1 to S11

Tables S1 to S10

References (*60-78*)

