## Supplementary Information for "CD36 homologs determine microbial resistance to the Lyme disease spirochete"

**This PDF includes:**

Materials and Methods
Figures S1 to S11
Tables S1 to S10

### Materials and Methods

Bacteria, mice, and ticks

*Escherichia coli* BL21 (DE3) cultures were grown at 37°C overnight in lysogeny broth (LB; Millipore Sigma) supplemented with 100 µg/ml ampicillin. Low passage *Borrelia burgdorferi* B31 clone MSK5 was grown in Barbour-Stoenner Kelly (BSK)-II medium with 6% normal rabbit serum (*60*). *B. burgdorferi* were grown at 34°C for tick cell stimulations and 37°C for mouse infections, never exceeding $10^8$ bacteria per ml. Plasmid profiling was performed, as described elsewhere (*60*). Age matched, six- to ten-week-old C3H/HeJ, and *Cd36*$^{-/-}$ mice were supplied by Jackson Laboratories. C57BL/6J mice were supplied by Jackson Laboratories or the University of Maryland Veterinary Resources. *I. scapularis* nymphs were supplied by Oklahoma State University and University of Minnesota breeding colonies. Ticks were housed upon arrival at 23°C with 85% relative humidity and a 12/10-hour light/dark photoperiod cycle. All mouse experiments were approved by the Institutional Biosafety (IBC, IBC-00002247) and Animal Care and Use (IACUC, #0119012) committees at the University of Maryland School of Medicine and complied with National Institutes of Health (NIH) guidelines (Office of Laboratory Animal Welfare [OLAW] assurance number A3200-01).

Tick cell culture

The *I. scapularis* cell lines IDE12 and ISE6 were cultured in L15C300 medium supplemented with 10% heat-inactivated fetal bovine serum (FBS, Millipore Sigma), 10% tryptose phosphate broth (BD), and 0.1% bovine lipoprotein concentrate (MP Biomedicals) at 34°C. Tick cells were grown to confluence and either seeded at $4x10^5$ cells/well in 24-well plates (Corning) or $1x10^6$ cells/well in 6-well plates (Millipore Sigma) or sub-cultured in capped T25 flasks (Greiner bio-one). All cell cultures were verified by PCR to be *Mycoplasma*-free (Southern Biotech).

*I. scapularis* Croquemort identification by mass spectrometry

Lipid binding partners were identified by performing co-immunoprecipitation with biotinylated POPG (Avanti). 50 μl of streptavidin beads were incubated with 50 μg biotinylated POPG in TBS buffer at 4°C for 1 hour. Residual biotin sites were blocked using biotin blocking buffer. 1 mg of ISE6 cell lysates were added and incubated overnight at 4°C. Proteins were eluted and measured by silver staining before performing mass spectrometry (University of Maryland School of Medicine Protein Analysis Laboratory). Identified proteins were filtered using the Contaminant Repository for Affinity Purification (CRAPome) database to remove common contaminants (*61*).

Plasmids

The plasmids pET-14b (#69660) and DsRed-N1 (a gift from Michael Davidson, #54493) were obtained from Novagen and Addgene, respectively. Constructs were amplified from ISE6 cell line complementary DNA (cDNA) with high-fidelity polymerase. For generating a His-tagged Croquemort ectodomain (Crq-His) recombinant protein, codon-optimized *I. scapularis croquemort* was cloned into the pET-14b vector with a primer set containing NdeI and XhoI restriction sites (table S9; pET14b-Crq-ecto-His). The ectodomain was predicted by using the TMHMM transmembrane helix prediction software to identify transmembrane domains (*22*). For ectopic expression of Crq-dsRed, full length *croquemort* was cloned into the DsRed-N1 vector with a primer set containing NheI and EcoRI restriction sites (table S9; DsRed-N1-Crq). All constructs were verified through Sanger sequencing.

Lipid analysis

Sample preparation was carried out at Metabolon Inc, in a manner similar to a previous study (*62*). Briefly, ISE6 cell samples were subjected to methanol extraction and split into aliquots for analysis by ultrahigh performance liquid chromatography/mass spectrometry (UHPLC/MS). The global biochemical profiling analysis comprised of four unique arms consisting of reverse phase chromatography positive ionization methods optimized for hydrophilic compounds (LC/MS Pos Polar) and hydrophobic compounds (LC/MS Pos Lipid), reverse phase chromatography with negative ionization conditions (LC/MS Neg), as well as a HILIC chromatography method coupled to negative (LC/MS Polar) (*63*). All methods alternated between full scan MS and data dependent $MS^n$ scans. The scan range varied slightly between methods but generally covered 70–1000 *m*/*z*.

Metabolites were identified by automated comparison of the ion features in the experimental samples to a reference library of chemical standard entries that included retention time, molecular weight (*m/z*), preferred adducts, and in-source fragments as well as associated MS spectra and curated by visual inspection for quality control using software developed at Metabolon. Identification of known chemical entities was based on comparison to metabolomic library entries of purified standards (*64*).

Recombinant Croquemort ectodomain (Crq-His) expression and purification.

His-tagged Croquemort ectodomain (Crq-His) was successfully purified using the *E. coli* BL21 expression system. To purify Crq-His, *E. coli* BL21 (DE3) were transformed with the pET-14b-Crq-His plasmid and streaked onto LB agar plates containing 100 ug/ml ampicillin (Quality Biological) at 37°C. Individual colonies were used to make starter cultures and were grown in antibiotic-containing LB broth at 37°C for 12-18 hours. Starter cultures were added to larger volumes of LB and grown until reaching an OD600 of 0.4-0.8. *E. coli* were induced with 0.5 mM isopropyl β-D-1-thiogalactopyranoside (IPTG) for 3 hours at 37°C. Cells were collected by centrifugation, resuspended in 1X PBS containing lysozyme, 0.05% CA630, and protease inhibitors (Thermo Scientific) and sonicated for 30 minutes using the Model 120 Sonic Dismembrator (Fisherbrand).

Samples were centrifuged at 12,000 rpm for 15 minutes. Pellets were resuspended in denaturation buffer (6 M guanidine-HCl, 300 mM NaCl, 20 mM Tris, pH 7.4) and incubated for 1 hour at 37°C. Supernatant was obtained and incubated with prepared Hi60 Ni Superflow resin (TakaraBio) for 1 hour at room temperature or overnight at 4°C. The resin was washed four times with denaturation buffer. To elute His-tagged protein, beads were incubated with elution buffer (6 M guanidine-HCl, 300 mM NaCl, 20 mM Tris, 150 mM imidazole, pH 7.4) for 15 minutes before eluting into 50% final concentration glycerol. Elution buffer was removed through two dialysis steps (first 6 hours, second overnight) against 1X PBS, 200 mM NaCl, 10% glycerol. Proteins were concentrated using Amicon Ultra centrifugal filters (Millipore Sigma) and quantified using bicinchoninic acid (BCA) assay (Thermo Scientific). For protein used in two-dimensional nuclear magnetic resonance spectroscopy (2D-NMR), individual colonies were used to inoculate MOPS minimal media containing just $^{15}NH_4Cl$ as the sole nitrogen source for the expression of $^{15}N$-labeled Crq-His. Cultures were induced and purified as described above.

<u>NMR experiments and sample preparation</u>

NMR samples contained 15 μM $^{15}N$-labeled Crq in PBS, 10% $D_2O$. A $^{1}H$-$^{15}N$ transverse relaxation optimized spectroscopy (TROSY) heteronuclear single-quantum coherence (HSQC) spectrum (*65*) was collected at 298 K on a Bruker Avance III 950 MHz spectrometer equipped with a *z*-gradient cryogenic probe. NMR data were processed with NMRPipe (*66*) and analyzed with CcpNmr Analysis (*67*). All proton chemical shifts were referenced to external trimethylsilyl propanoic acid (TSP) at 25 °C (0.00 ppm) with respect to residual $H_2O$ (4.698 ppm). $^{1}H$-$^{15}N$ chemical shifts were indirectly referenced using a zero-point frequency ratio of 0.101329118.

Surface plasmon resonance

Surface plasmon resonance (SPR) of protein-lipid interactions was performed using a Biacore T200 instrument (GE Healthcare) (University of Maryland School of Medicine Biosensor Core). The carboxymethyl-dextran surface of a Biacore CM5 chip (flow cell-2, GE Healthcare) was activated with a 35 µl injection of 0.1 M NHS and 0.1 M EDC in water. Crq-His or a negative control protein (100 µl of 10 µg/ml dilution) were directly immobilized to the chip (9200 resonance units). Remaining NHS-ester sites in the flow cell were blocked with 1M ethanolamine, pH 8.2. The flow cell-1 of the same CM5 chip was used as reference and activated in a similar manner. Analytes (POPG) were diluted in HBS-EP buffer with 0.05% P20 (Activa) and injected into the flow cells to calculate association. The surface was washed with buffer to determine the dissociation of analyte-ligand complexes. Data was analyzed using the BIAeval 2.0 software and dissociation constants were calculated ($K_D = K_d/K_a$). Reference surface data (flow cell-1) were subtracted from the reaction surface data to eliminate refractive-index changes of the solution, injection noise, and non-specific binding to the blank surface. Furthermore, the signal from a blank injection with buffer alone was subtracted from the resulting reaction surface data.

RNA interference and tick experiments

For *in vitro* experiments, small interfering RNAs (siRNA) for *croquemort*, *relish*, *dorsal*, and *tak1* and their scrambled controls (scRNA) were synthesized by Millipore Sigma with dTdT overhangs. IDE12 cells were plated at $4 \times 10^5$ cells per well (24 well plate, RNA extraction) or $1 \times 10^6$ cells per well (6 well plate, protein extraction). siRNAs (1 µg per ml) were transfected into IDE12 cells using Lipofectamine 3000 (ThermoFisher). After 7 days, cells were either harvested without stimulation or stimulated with POPG before harvesting. For protein extraction, IDE12 cells were harvested, washed with PBS, resuspended in 1X RIPA lysis buffer (Millipore Sigma) with Halt protease and phosphatase inhibitor cocktails (Thermo Scientific), and stored at -80°C.

For RNA extraction, IDE12 cells were harvested in Trizol (Ambion) and stored at -80°C.

For *in vivo* experiments, siRNAs for *croquemort*, *relish*, *jun*, and *dorsal* and their scrambled controls were synthesized using the Silencer siRNA construction kit (Thermo Scientific) using the primers in table S9. *I. scapularis* nymphs were microinjected with 20 – 40 ng of siRNA or scRNA. Ticks rested overnight before being placed on anesthetized C3H mice for 20 minutes. Ticks were allowed to feed up to five days unless otherwise stated. Fully engorged nymphs were collected in tubes, weighed, and either placed in a humidified chamber for molting or frozen at -80°C. Tubes containing single ticks were submerged in liquid nitrogen and homogenized prior to RNA extraction or methanol extraction. Tick feeding time courses were performed with uninfected male C57BL/6J mice.

Mouse infections

C3H mice were used for *B. burgdorferi* tick acquisition experiments. Briefly, *B. burgdorferi* were washed and resuspended in 50% 1X PBS, 50% normal rabbit serum at a concentration of $1x10^6$ *B. burgdorferi* per ml. Mice were anesthetized with a ketamine-xylazine sedative and an area of ~2 $cm^2$ was shaved on the back using an electric clipper. Mice were injected intradermally with 100 µl of inoculum ($1x10^5$ total *B. burgdorferi*). Mice were maintained for at least 14 days prior to tick placement.

Macrophage differentiation and immune measurements

Bone marrow-derived macrophages (BMDMs) were generated from C57BL/6J and *Cd36*$^{-/-}$ mice. Briefly, mice were euthanized via $CO_2$ and femurs were dissected. Bone marrow was flushed by injecting differentiating medium consisting of DMEM supplemented with 30% L929 conditioned medium, 10% FBS (Gemini Bio-products), 1X Amphotericin B (Gibco), and 1X penicillin/streptomycin (Corning) into one end of the femur with a 25-gauge needle. Cells were seeded in 90 mm Petri dish plates and incubated at 37°C 5% $CO_2$. Cells were incubated for 7 days until complete differentiation and replated into 24 well plates at a density of $1x10^6$ cells per well. For microbial stimulations, cells were stimulated with *B. burgdorferi* (MOI 50) for indicated time points. Cell pellets were resuspended in 1X RIPA lysis buffer (Millipore Sigma) containing 1X Halt Protease and Phosphatase Inhibitor Cocktail (Thermo Scientific) and processed for western blotting. Supernatants were collected for cytokine quantification and analyzed by ELISA. Cytokine concentrations were measured using the IL-6 Mouse ELISA Kit (Invitrogen), KC/CXCL1 Mouse ELISA Kit (Invitrogen), and Mouse IL-1 β/IL-1F2 Quantikine Kit (R&D Systems), as per the manufacturers' instructions.

Western blotting

Western blotting was performed as previously described *(11).* Briefly, protein lysates were quantified by BCA assay (Thermo Scientific). Equal amounts of protein were boiled in 6X Laemmli sample buffer (Alfa Aesar) containing 5% β-mercaptoethanol. Samples were loaded in mini-PROTEAN gels (Biorad) subjected to sodium dodecyl sulfate polyacrylamide gel electrophoresis (SDS-PAGE). Proteins were then transferred onto PVDF membranes (Biorad) and membranes were blocked with 5% milk (Biorad) for 1 hour in PBS-T. Primary antibodies were incubated overnight at 4°C in PBS-T with 3% bovine serum albumin (BSA). Blots were washed four times in PBS-T and incubated with secondary antibodies for at least 1 hour at room temperature with rocking. Blots were washed four times in PBS-T, incubated with enhanced chemiluminescence (ECL) substrate solution for 1 minute (Thermo Scientific), and imaged.

Quantitative reverse transcription polymerase chain reaction (qRT-PCR).

The PureLink RNA Mini kit (Invitrogen) was used to extract RNA from cells or engorged ticks preserved in Trizol. cDNA was synthesized with the Verso cDNA Synthesis Kit (ThermoFisher). qRT-PCR was performed with the CFX96 Touch Real-Time PCR Detection

System (Biorad). Levels of *B. burgdorferi* gene expression were measured by absolute quantification and normalized to tick *actin* (for tick experiments). Levels of expression for all other genes were measured by relative quantification. Transcriptional silencing and *B. burgdorferi* burden were quantified using the primers listed in table S9.

Protein alignments and modeling.

Amino acids 30 - 440 of *I.s.* Croquemort were used for ectodomain modeling based on TMHMM software predictions and previous work on CD36 protein family structure (*22*). The Croquemort ectodomain was modeled to the crystal structures of human LIMP-2 (PDB:4F7b) (*20*) and human CD36 (PDB:5LGD) (*21*) using Protein Homology/Analogy Recognition Engine (Phyre) 2 (*24*). Predicted interactions between POPG and the Croquemort Phyre2 model were subsequently measured using AutoDock (*26*). Additional structural predictions were performed using the AlphaFold source code (*25*) and accessed through NMRbox (*68*). Root-mean-square deviations (RMSDs) were calculated in Pymol. Sequence alignments were generated using Multiple Sequence Comparison by Log Expectation (MUSCLE) (*69*) and visualized using JalView (*70*). Conserved residues were identified from sequence alignments with LIMP-2 (Q14108) and CD36 (P16671).

Subcellular fractionation.

Subcellular fractionation was performed as previously described (*71*). Briefly, ISE6 cells were resuspended in fractionation buffer and passed through a 27-gauge needle 10 times. Cells were incubated on ice for 20 minutes and then centrifuged at 10,000 x g for 5 minutes. To isolate the membrane fraction, the supernatant was transferred and centrifuged at 100,000 x g for 1 hour. The supernatant was reserved (cytosol). The pellet was washed in 400 µl, passed through a 25-gauge needle, and re-centrifuged at 100,000 x g for 45 minutes. The remaining pellet (membrane) was resuspended in TBS containing 0.1% sodium dodecyl sulfate (SDS).

Nucleofection and ectopic expression of Crq-dsRed

800 ng of Crq-DsRed-N1 construct was nucleofected in $1x10^6$ tick cells (ISE6 and IDE12) using the 4D-Nucleofector System (Lonza Bioscience). $1x10^6$ tick cells were centrifuged at 100 x g for 10 minutes to pellet the cells. The pellet was washed with 10 ml PBS, resuspended in 20 µl SF buffer (Lonza Bioscience) and 800 ng of Crq-DsRed-N1 construct was added to the cell suspension. The nucleofection mix was added to a multiwell cuvette, inserted into the nucleofector and pulsed using pulse condition EN150. The cells were allowed to sit in the cuvette for 10 minutes post-nucleofection before being added to pre-warmed L15C300 complete media in a 12 well plate. After 72 hours, cells were fixed with 4% paraformaldehyde (PFA), stained with CellMask green plasma membrane (Invitrogen) and Hoechst nuclear (AAT Bioquest) stains, and observed under the fluorescence microscope.

Immunostaining and confocal microscopy.

IDE12 cells were spun down onto microscope slides using a Cytospin (Thermo Scientific) and fixed with 4% PFA. Cells were washed three times with PBS and blocked with 3% bovine serum albumin (BSA) in PBS for 1 hour at room temperature. Primary antibody or rabbit IgG isotype control was prepared in blocking buffer and incubated with cells overnight at 4°C. The following day, slides were washed three times with PBS before addition of goat anti-rabbit IgG secondary antibody (Invitrogen). Slides were washed three times with PBS and SlowFade Gold Antifade mountant containing DAPI (Invitrogen) was added. Coverslips were mounted and sealed with nail polish.

Microscopy images were obtained using a Nikon W-1 spinning disk confocal microscope. The following laser channels were used for imaging Crq-DsRed-N1 cells: 561 nm (Crq-DsRed), 488 nm (GFP, plasma membrane), and 405 nm (DAPI). An excitation wavelength of 558 nm and an emission wavelength of 583 nm were used for obtaining images of Crq-DsRed-N1. For immunostaining, the following laser channels were used: 640 nm (mCherry, anti-rabbit Alexa Fluor 594), and 405 nm (DAPI). An excitation wavelength of 590 nm and an emission wavelength of 617 nm were used for obtaining images of IDE12 cells stained with anti-Crq primary antibody.

Electrophoretic mobility shift assay (EMSA)

The putative NF-kB binding site to the *Ixodes scapularis ctenidin-1* promoter (XM_029986701.1, NCBI *Ixodes scapularis* Annotation Release 100) was inferred using two free software: LASAGNA-Search v2.0 (*72*) and Promo v3.0.2 (*73*). A consensus region between each software was identify at position -134-116, corresponding to the sequence 5' CCTAGGCGTCTCCCCGGC 3', which was used to synthesize 5' biotinylated or unbiotinylated double-stranded DNA probes (Thermo Scientific) for the EMSA. Nuclear extracts (NE) from $3x10^6$ IDE12 cells stimulated with 10 ng/ml POPG were obtained using NE-PER Nuclear and Cytoplasmic Extraction Reagents (Thermo Scientific), following the manufacturer's instructions, and stored at -80C until use. Binding reactions were performed according to the LightShift™ Chemiluminescent EMSA Kit instructions (Thermo Scientific). Briefly, binding reactions contained 1X binding buffer, 50 ng of poly(dI-dC), 100 fm of biotinylated probes and 3 µl (~10 µg) of NE in a final volume of 20 µl. For competition assays, 1 pm and 20 pm of unlabeled probes (10:1 and 200:1 unlabeled/label probe ratio, respectively) were included in the mix. To test the specificity of the transcription factor that binds to the probe, 0.5 µg, 1.5 µg and 5 µg of mouse antiserum containing *I. scapularis* N-Rel antibodies (Genscript) were added to the mix. For those reactions, 5 µg of mouse (G3A1) mAb IgG1 isotype control (Cell Signaling) was used as control. After 20 minutes of incubation at room temperature, the DNA probe-protein complex interactions were resolved by electrophoresis on a 5% Mini-PROTEAN® TBE Gel (Bio-Rad). Subsequently, the gel was transferred to a Biodyne B Precut Nylon Membrane (Thermo Scientific), and DNA was crosslinked to the membrane using an UV-light transilluminator for 7 min. Finally, the membrane was developed per the manufacturer's protocol.

Ecdysteroid quantification.

Fully repleted, whole ticks were frozen in liquid nitrogen and thoroughly homogenized using pestles. 500 μl of methanol was added to homogenate and samples were centrifuged at 10,000 x g for 5 minutes. The supernatants were collected on ice and the remaining pellet was re-extracted by an additional 500 μl of methanol. The samples were dried using a SpeedVac (Thermo Scientific) and dissolved in EIA buffer (Cayman Chemical). Ecdysteroid concentrations were determined using a 20-hydroxyecdysone EIA kit (Cayman Chemical).

Biobank study population

The study was approved by the Institutional Review Board of the Icahn School of Medicine at Mount Sinai (GCO#07-0529; STUDY-11-01139) according to the tenets of the Declaration of Helsinki. Informed consent was obtained for all participants for the storage of biological specimens, genetic sequencing, and access to de-identified electronic health record data (*e.g.,* clinical diagnoses, laboratory results, family history of disease, medical history, medications, demographics). All study participants in Bio*Me* were recruited from outpatient sites in the Mount Sinai Health System in New York City and the first 31,250 participants underwent exome sequencing. Quality control was applied in which samples with discordance between genetic and recorded sex, low coverage, contamination, or duplicate samples were removed. In addition, samples without complete demographic data, younger than 20 years, related up to third degree relatives, or missing ICD-10 data were excluded to produce the final study population.

Clinical outcomes

Lyme disease was the primary outcome and cases were identified by the presence of a physician-documented International Classification of Diseases-Clinical Modification 10 (ICD-10) diagnosis code of A69.2 or a laboratory-confirmed result. The latter was defined by a positive IgG/IgM *B. burgdorferi* antibody test and/or positive *B. burgdorferi* PCR result in accordance with the Centers for Disease Control and Prevention guidelines. Secondary outcomes comprised immunological laboratory measurements, for which median values were obtained: percent neutrophils (n=24,296 individuals), percent lymphocytes (n=24,335), complement C4 (n=2,111), IL-1β (n=93), and IL-8 (n=93). Median total cholesterol level and statin usage were also ascertained for all participants. Other clinical variables (age, sex, BMI, etc.) were taken at baseline when the participants were first enrolled at BioMe.

Exome sequencing and identification of *CD36* LoF variants

Exome sequencing and quality control are described extensively elsewhere (*46*). Briefly, samples were prepared, and variant call files (VCFs) were generated with Illumina v4 HiSeq 2500 by the Regeneron Genetics Center. These were subjected to the Goldilocks Filter (GF) (*74*) and standard quality control procedures were applied for depth-normalized quality scores, depth of coverage, allele balance, and missing genotypes. We identified five LoF variants in *CD36* based on annotations of frameshift, splice donor, splice acceptor, stop gained, stop lost, or start lost using Variant Effect Predictor (VEP) (*75*). Three rare *CD36* LoF variants with allele frequency of less than or approximately equal to 1% were obtained based on a similar threshold previously used to enrich for rare, highly impactful variants (*76*) and used for all downstream analyses.

Association testing of *CD36* LoF burden with clinical outcomes

We aggregated all alleles for rare *CD36* LoF variants carried by each individual into a single burden score and assessed this *CD36* LoF burden for association with Lyme disease. Logistic regression with Firth's penalized likelihood was used, adjusted for age, sex, BMI, and 10 genetic PCs. Firth's approach offsets the first-order term in the asymptotic expansion of the bias of small sample sizes when estimating the maximum likelihood (*77*). *CD36* LoF burden was also evaluated for association with quantitative immune measurements using linear regression adjusted for the aforementioned covariates.

Resources and reagents

Additional information provided in table S10.

Statistical analysis

Statistical significance was assessed with the Student's t test and unpaired *t* test with Welch's correction. Two-way ANOVA post-hoc Sidak test for multiple comparisons was also used whenever appropriate. Molting curves were analyzed with the Log-rank (Mantel-Cox) test. We used GraphPad PRISM® (GraphPad Software version 9.1.0) for all statistical analyses. Outliers were detected by a Graphpad Quickcals program (https://www.graphpad.com/quickcalcs/Grubbs1.cfm

### Supplementary Figures

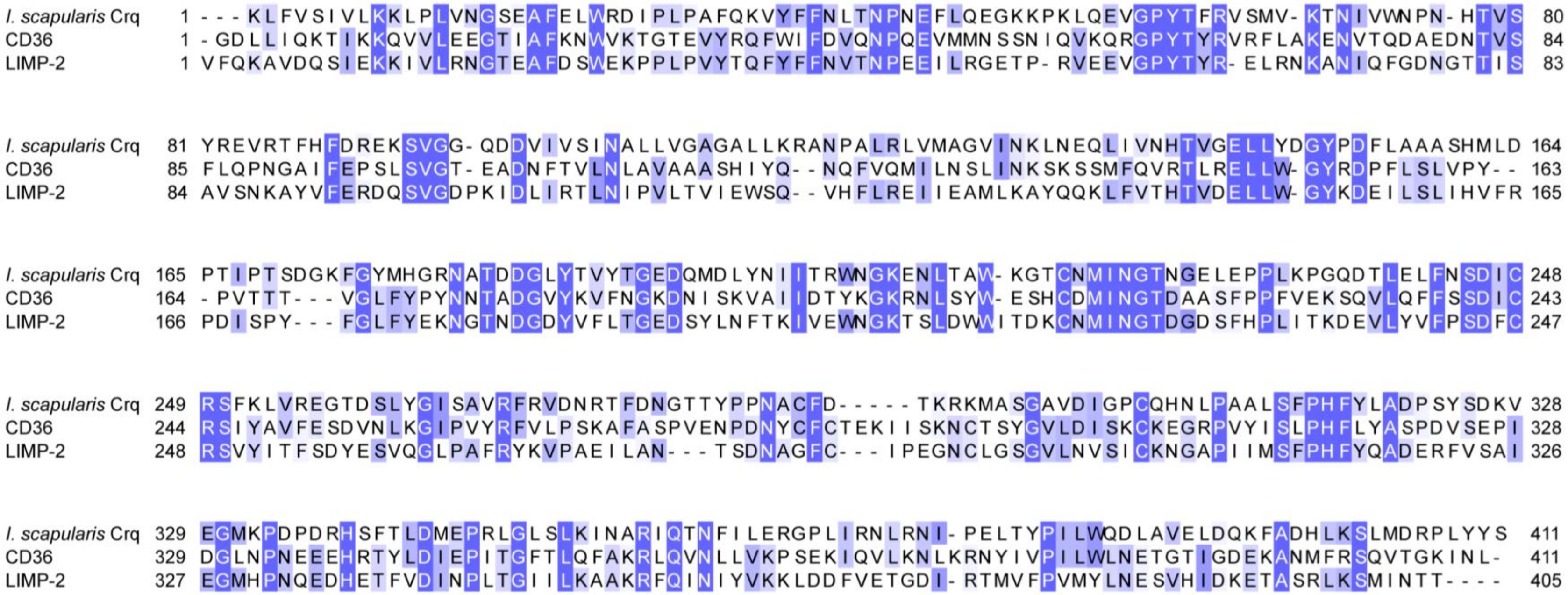

**Fig. S1. Protein alignment of the *I. scapularis* Croquemort (Crq), CD36, and LIMP-2 ectodomains.** The sequence of *I. scapularis* Crq was empirically determined from ISE6 cells and translated into the amino acid sequence. The Crq ectodomain (amino acids 30-440) was identified from TMHMM predictions and aligned to the CD36 (amino acids 30-440) and LIMP-2 (amino acids 28-432) ectodomains. The sequence alignment was generated using Multiple Sequence Comparison by Log Expectation (MUSCLE) and visualized with JalView. Residues are color-coded by the conservation index and percentage identity to *I. scapularis* Crq.

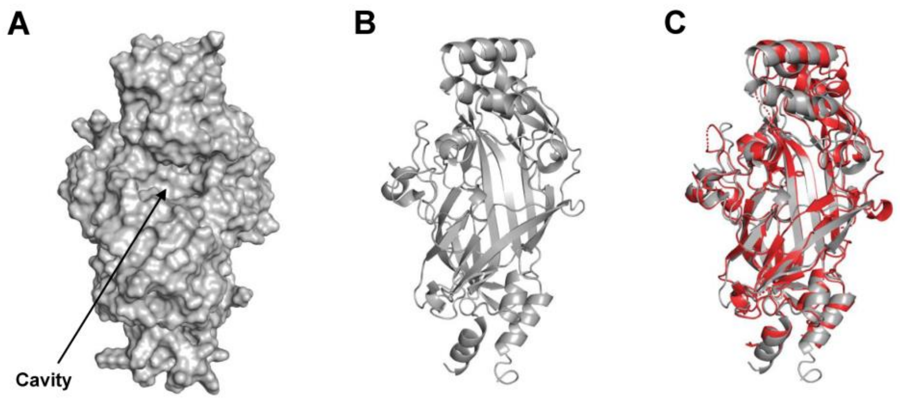

**Fig. S2. Predicted structure of the Crq ectodomain using AlphaFold.** The Crq ectodomain as predicted by AlphaFold shown as a surface representation (**A**) and a Ribbon diagram (**B**). **C.** The predicted structure of Crq (grey) aligned to the Phyre2 generated model of Crq (red) (RMSD = 0.89 Å) showing structural similarity between the models. Structures were visualized in Pymol. RMSD=root-mean-square deviation.

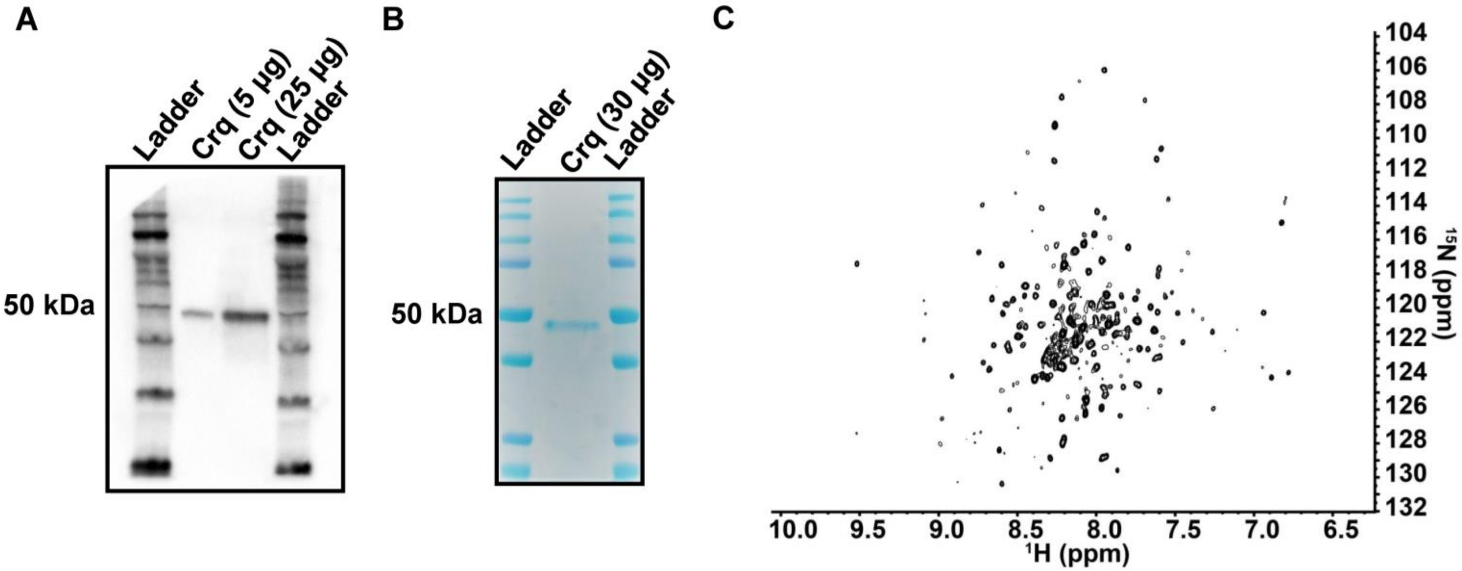

**Fig. S3. Protein purification and 2D-nuclear magnetic resonance spectroscopy of Crq-His. A.** Representative western blot of purified Crq-His (anti-His). **B.** Coomassie blue staining of purified Crq-His. **C.** 2D-[$^{1}$H, $^{15}$N]-TROSY-HSQC spectrum of Crq-His at 950 MHz ($^{1}$H dimension). Approximately 260 of the 384 possible correlations were observed in the 2D HSQC NMR spectrum. Correlations were well dispersed in both the $^{1}$H and $^{15}$N dimensions, as compared to chemical shift values in an unfolded protein. Correlations were consistent with $H_N$ moieties existing in folded regions of the Crq-His protein. That ~30% of the possible $H_N$ correlations were not observed suggest that at least some region(s) of Crq-His were flexible on the chemical shift timescale (*78*). TROSY=Transverse relaxation optimized spectroscopy; HSQC=Heteronuclear Single Quantum Coherence; NMR=Nuclear magnetic resonance.

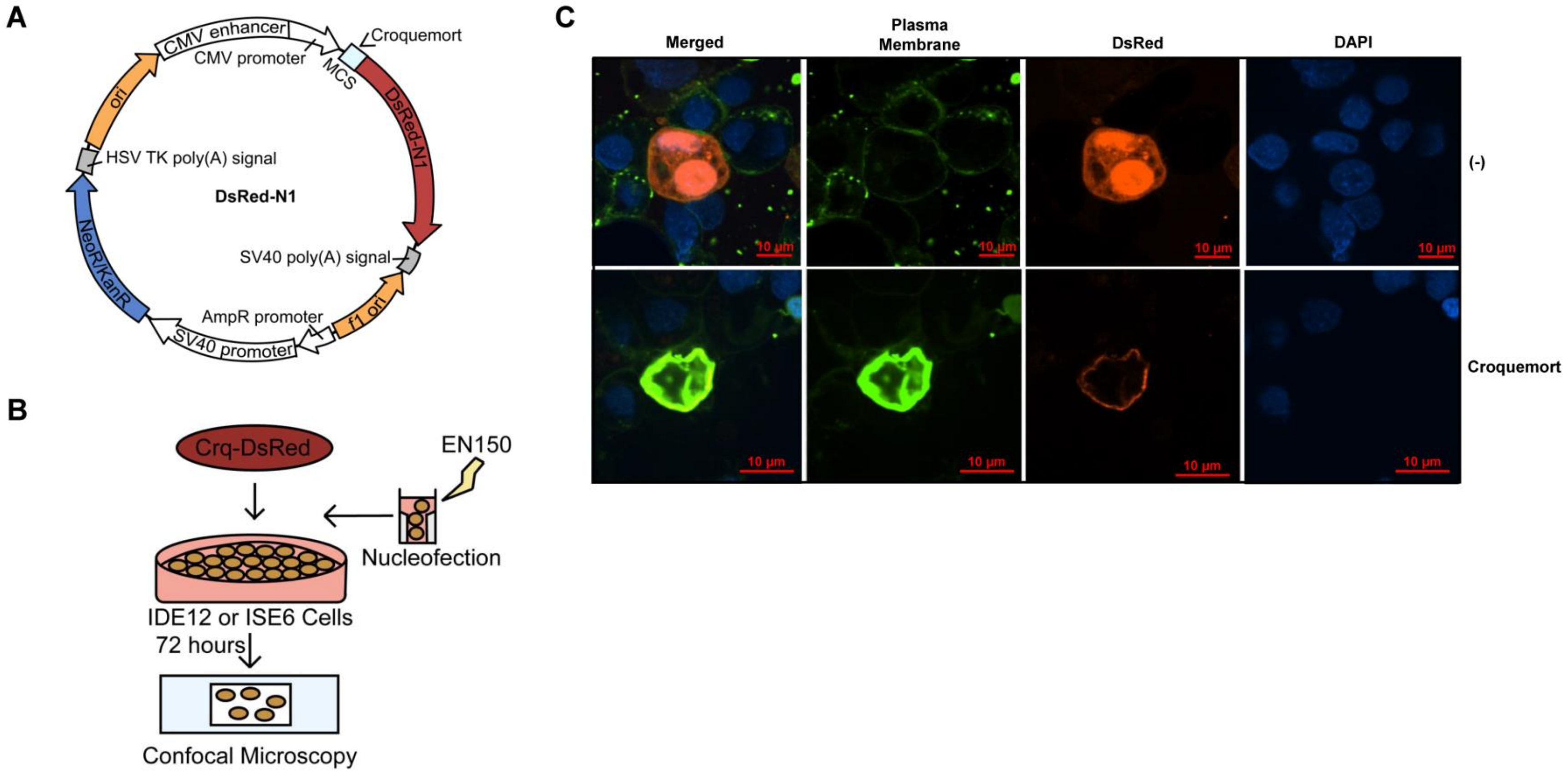

**Fig. S4. Confocal microscopy of Crq in ISE6 cells. A.** Diagram of the Crq-DsRed-N1 construct used for ectopic expression. **B.** Schematic of ectopic expression workflow. IDE12 or ISE6 cells were nucleofected (SF buffer, pulse condition EN150) with plasmid containing Crq-DsRed or empty vector. Cells were plated and incubated for 72 hours prior to confocal imaging. **C.** Ectopic expression of DsRed-tagged Crq in the ISE6 tick cell line. Green - plasma membrane; blue – DAPI (4',6-diamidino-2-phenylindole). Data represent one of two independent experiments. (-)=empty vector; ori=origin of replication; MCS=multiple cloning site;

NeoR=neomycin resistance; KanR=kanamycin resistance.

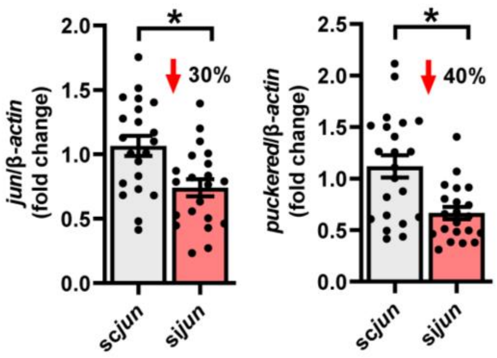

**Fig. S5. *puckered* expression in *jun* silenced ticks.** Ticks were microinjected with *jun* siRNA (si*jun*) or scrambled RNA (sc*jun*) and fed on *B. burgdorferi*-infected mice until repletion (*n*=21). Results are represented as means +/- SE. Statistical significance was evaluated by an unpaired *t* test with Welch's correction. *, *p*<0.05.

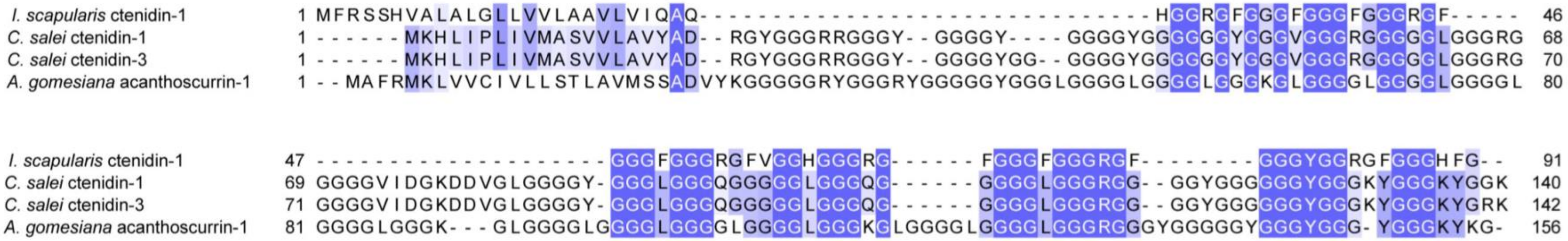

**Fig. S6. Peptide alignment of *I. scapularis* ctenidin-1 and related glycine-rich peptides from spiders.** The sequence of an *I. scapularis* ctenidin-1 peptide (annotated as "ctenidin-1-like") was aligned to the *Cupiennius salei* (tiger spider) ctenidin-1, *C. salei* ctenidin-3, and *Acanthoscurria gomesiana* (tarantula) acanthoscurrin-1 peptide sequences. The sequence alignment was generated using Multiple Sequence Comparison by Log Expectation (MUSCLE) and visualized with JalView. Residues are color-coded by the conservation index and percentage identity to *I. scapularis* ctenidin-1.

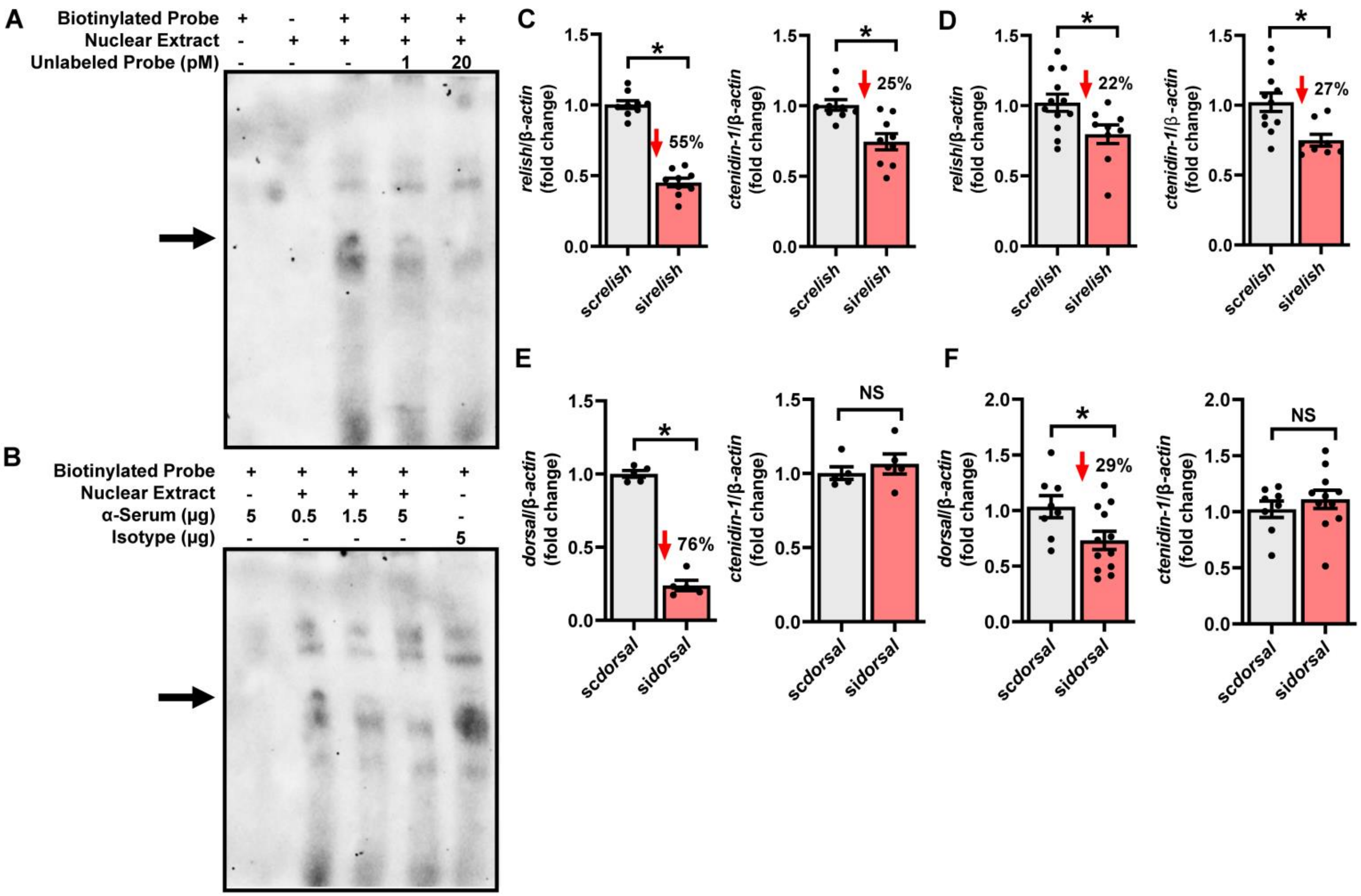

**Fig. S7. *ctenidin-1* is regulated by Relish, but not Dorsal. A.** Electrophoretic mobility shift assay (EMSA) of IDE12 cell nuclear extract, biotinylated *ctenidin-1* promoter probe, and unlabeled probe. Nuclear extract was incubated with biotinylated promoter probe and increasing concentrations of unlabeled probe. Band appears when biotinylated probe was added to nuclear extract, which disappears following addition of unlabeled probe in a dose-dependent manner. Arrows denote band of interest. **B.** EMSA of IDE12 cell nuclear extract, biotinylated *ctenidin-1* promoter probe, and mouse antiserum containing antibodies for *I. scapularis* N-Rel. Nuclear extract was incubated with biotinylated promoter probe and increasing concentrations of N-Rel antiserum. Specific band (as in fig. S7A) disappeared following addition of N-Rel antiserum in a dose-dependent manner. Arrows denote band of interest. **C.** *ctenidin-1* expression and *relish* silencing efficiency in IDE12 cells. Cells were transfected with *relish* siRNA (si*relish*) or scrambled RNA (sc*relish*) (*n*=9). Two biological replicates were performed. **D.** *ctenidin-1* expression and *relish* silencing efficiency in ticks microinjected with si*relish* or sc*relish*. Ticks were allowed to feed until repletion (*n*=8-12). **E.** *ctenidin-1* expression and *dorsal* silencing efficiency in IDE12 cells. Cells were transfected with dorsal siRNA (si*dorsal*) or scrambled RNA (sc*dorsal*) (*n*=5). Two biological replicates were performed. **F.** *ctenidin-1* expression and *dorsal* silencing efficiency in ticks microinjected with si*dorsal* or sc*dorsal*. Ticks were allowed to feed until repletion (*n*=8-12). Results are represented as means +/- SE. Statistical significance was evaluated by an unpaired *t* test with Welch's correction. *, $p<0.05$.

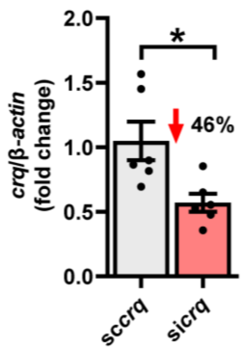

**Fig. S8. *crq* silencing efficiency in IDE12 cells.** Silencing efficiency of *crq* siRNA used in Figure 2L. Cells were transfected with *crq* siRNA (si*crq*) or scrambled RNA (sc*crq*) (*n*=6). Results are represented as means +/- SE. Data represent one of two independent experiments. Statistical significance was evaluated by an unpaired *t* test with Welch's correction. *, $p$<0.05.

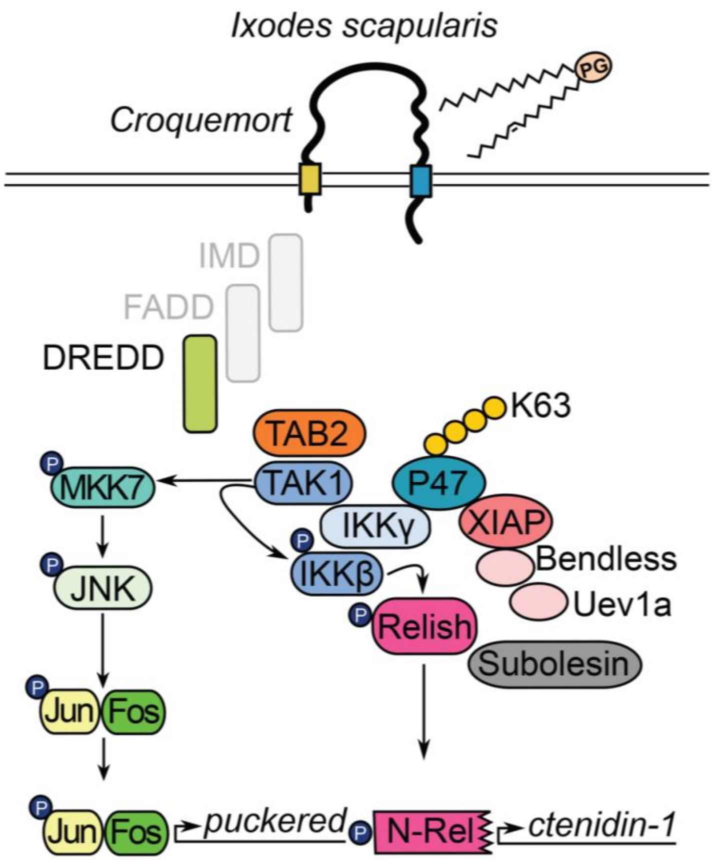

**Fig. S9. Diagram of signaling model in *I. scapularis*.** *I. scapularis* Croquemort (Crq) binds to POPG. Following POPG stimulation, Crq relays an antimicrobial program through the protein TAK1, which subsequently activates immune deficiency (IMD) and JNK signaling components. The IMD-specific transcription factor Relish promotes expression of the constitutively expressed antimicrobial peptide *ctenidin-1*, while the transcription factor Jun promotes *puckered* expression. Crq utilizes both the IMD and JNK pathways to restrict colonization by *B. burgdorferi*.

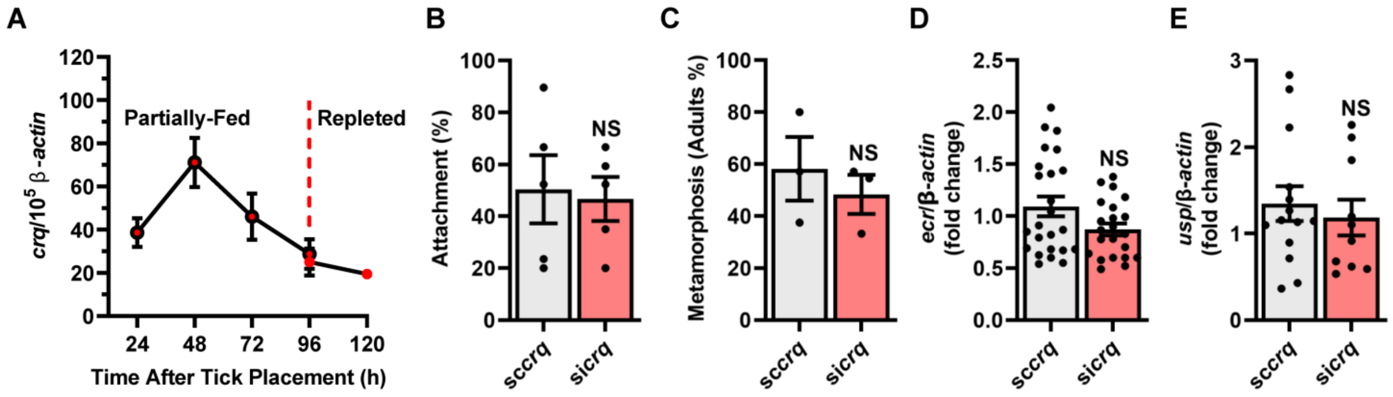

**Fig. S10. Fitness parameters in *crq* silenced nymphs. A.** Expression of *crq* over time of feeding in naïve *I. scapularis* ticks (*n*=3-6 per time point, 3 nymphs pooled per replicate). Results are displayed as copies of *crq* per $10^5$ copies of *I. scapularis actin*. **B.** Attachment of ticks microinjected with *crq* siRNA (si*crq*) or scrambled RNA (sc*crq*). ~25 ticks were placed per mouse and allowed to feed for up to five days. Each point represents the percentage of fully repleted ticks recovered from a mouse out of total placed (*n*=5). **C.** Metamorphosis of si*crq* and sc*crq* ticks. Ticks were allowed to feed until repletion and subsequently monitored for molting. Each point represents the percentage of ticks that molted out of total recovered per mouse (*n*=3). **D** and **E.** Expression of ecdysteroid receptor genes *ecr* (**D**) and *usp* (**E**) in fully repleted si*crq* and sc*crq* ticks (*n*=10-24). Results are represented as means +/- SE. Statistical significance was evaluated by an unpaired *t* test with Welch's correction. NS=not significant.

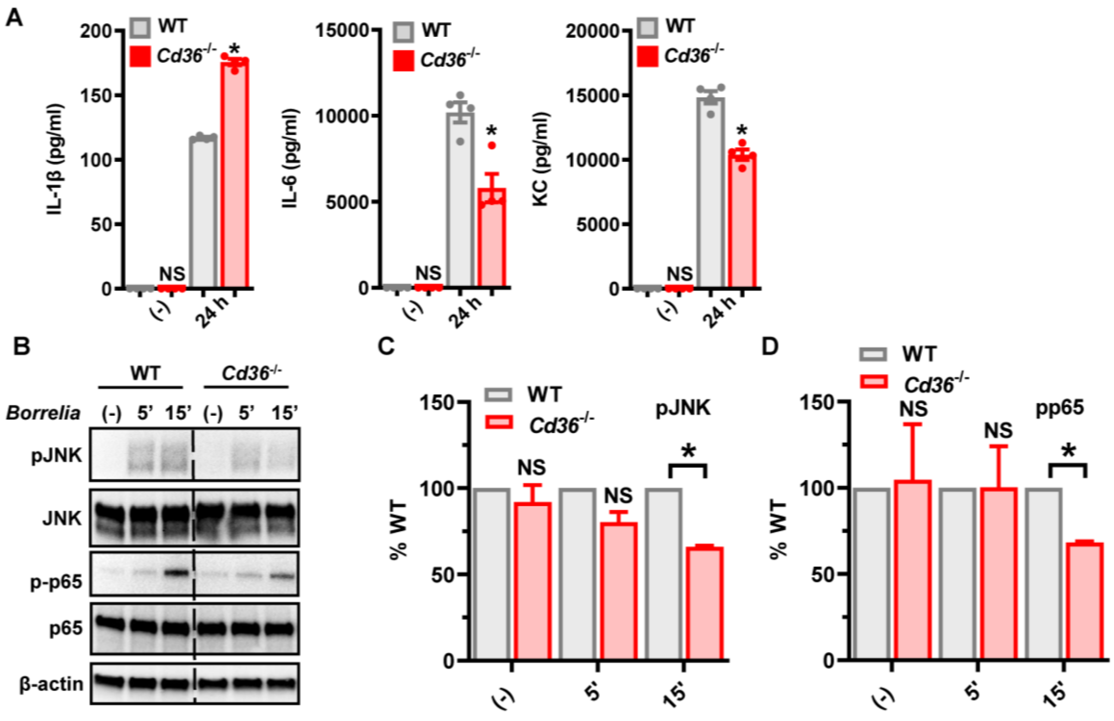

**Fig. S11. Immune profiling of murine *Cd36*$^{-/-}$ bone marrow-derived macrophages (BMDMs). A.** Proinflammatory cytokine measurements of BMDMs isolated from wild-type (WT) C57BL/6 and *Cd36*$^{-/-}$ mice. $1x10^6$ of BMDMs were stimulated with *B. burgdorferi* (MOI 50) for 24 hours. Cytokines were detected by enzyme linked immunoassay (ELISA) (*n*=2-4). **B-D.** WT and *Cd36*$^{-/-}$ BMDMs ($1x10^6$) were stimulated with *B. burgdorferi* (MOI 50) for indicated time points and analyzed for JNK and NF-κB (p65) activation. Normalized data for *Cd36*$^{-/-}$ were divided by corresponding control values and expressed as a percentage of WT. pJNK values are normalized to JNK (p52) and pp65 values are normalized to p65. Western blot bands were quantified using ImageJ. Western blot images show one representative experiment (*n*=2). Results are represented as means +/- SE. Statistical significance was evaluated by an unpaired *t* test with Welch’s correction (A) or a Student’s t test (C, D). (-)=unstimulated; h=hours; pg=picograms; ml=milliliter; IL=interleukin, KC=keratinocytes-derived chemokine; WT=wild type; pJNK=phospho-JNK; pp65=phospho-p65; *, $p<0.05$, NS=not significant.

### Supplementary Tables

Table S1. Lipid profile for *B. burgdorferi*-stimulated ISE6 tick cells.

| Lipid/Pathway | Biochemical Name | 1 h | 24 h |
|---|---|---|---|
| Phosphatidylcholine | 1-myristoyl-2-palmitoyl-GPC (14:0/16:0) | 4.2 | 10.71 |
| | 1,2-dipalmitoyl-GPC (16:0/16:0) | 27.63 | 27.09 |
| | 1-palmitoyl-2-palmitoleoyl-GPC (16:0/16:1) | 1.05 | 1.23 |
| | 1-palmitoyl-2-docosahexaenoyl-GPC (16:0/22:6) | 0.68 | 0.83 |
| Phosphatidylethanolamine | 1-oleoyl-2-linoleoyl-GPE (18:1/18:2) | 0.85 | 0.94 |
| | 1-oleoyl-2-arachidonoyl-GPE (18:1/20:4) | 0.86 | 0.86 |
| Phosphatidylglycerol | **1-palmitoyl-2-oleoyl-*sn*-glycero-phosphoglycerol (16:0/18:1)** | **14.72** | **12.25** |
| | 1-stearoyl-2-oleoyl-GPG (18:0/18:1) | 2.25 | 3.34 |
| Lysophospholipid | 1-palmitoyl-GPC (16:0) | 0.53 | 0.72 |
| | 1-palmitoleoyl-GPC (16:1) | 0.37 | 0.43 |
| | 2-palmitoleoyl-GPC (16:1) | 0.52 | 0.65 |
| | 1-stearoyl-GPC (18:0) | 0.6 | 0.78 |
| | 1-oleoyl-GPC (18:1) | 0.39 | 0.46 |
| | 1-linoleoyl-GPC (18:2) | 0.48 | 0.49 |
| | 1-palmitoyl-GPE (16:0) | 0.54 | 0.57 |
| | 1-oleoyl-GPE (18:1) | 0.4 | 0.41 |
| | 1-linoleoyl-GPE (18:2) | 0.34 | 0.38 |
| | 1-stearoyl-GPS (18:0) | 0.58 | 0.6 |
| | 1-oleoyl-GPS (18:1) | 0.27 | 0.29 |
| | 1-palmitoyl-GPG (16:0) | 2.51 | 2.42 |
| | 1-stearoyl-GPG (18:0) | 2.09 | 4.47 |
| | 1-oleoyl-GPG (18:1) | 1.88 | 2.58 |
| Glycerolipid Metabolism | glycerol | 0.05 | 0.05 |
| | glycerophosphoglycerol | 2.7 | 3.75 |
| Diacylglycerol | palmitoyl-oleoyl-glycerol (16:0/18:1) | 1.84 | 1.02 |
| Galactosyl Glycerolipids | 1-palmitoyl-2-linoleoyl-galactosylglycerol (16:0/18:2) | 4.64 | 3.96 |
| Sterol | cholesterol | 0.89 | 0.83 |

Table S2. Proteomics results for pulldown assay with biotinylated POPG.

| Entry | Protein | Gene | Length | Score | Coverage | # Proteins | # Unique Peptides | # Peptides | # PSMs | # AAs | MW [KDa] | Calculated pI |
|---|---|---|---|---|---|---|---|---|---|---|---|---|
| B7QEZ9 | Translocon-associated protein subunit β (TRAP-β) (Signal sequence receptor subunit β) | 8040271 IscW_ISCW012704 | 191 | 22.12 | 30.37 | 1 | 3 | 3 | 6 | 191 | 21.4 | 7.4 |
| B7PYE7 | Endoplasmic reticulum transmembrane protein | 8033203 IscW_ISCW009367 | 243 | 20.26 | 23.46 | 1 | 4 | 4 | 7 | 243 | 27.7 | 9.04 |
| B7PN29 | Steroid membrane receptor Hpr6.6/25-Dx, putative (EC 1.3.1.74) | 8050754 IscW_ISCW018528 | 214 | 14.96 | 29.91 | 1 | 5 | 5 | 7 | 214 | 23.8 | 4.89 |
| B7Q926 | Signal sequence receptor subunit δ (Translocon-associated protein subunit δ) | 8037713 IscW_ISCW010675 | 173 | 13.47 | 27.75 | 1 | 4 | 4 | 4 | 173 | 19.1 | 6.52 |
| B7PQK7 | Integrin α-ps, putative (EC 3.1.4.50) | 8052228 IscW_ISCW005672 | 1056 | 12.58 | 8.9 | 1 | 8 | 8 | 8 | 1056 | 117.3 | 6.44 |
| B7PDC8 | Integrin α-ps, putative (EC 3.1.4.50) | 8028238 IscW_ISCW003187 | 565 | 6.02 | 4.96 | 1 | 2 | 2 | 2 | 565 | 61.9 | 5.81 |
| B7QK99 | Signal recognition particle receptor subunit β | 8042520 IscW_ISCW013711 | 249 | 5.41 | 11.24 | 1 | 2 | 2 | 2 | 249 | 27.5 | 9.79 |
| **B7PC41** | **Scavenger receptor class B type I** | **8027712 IscW_ISCW002412** | **351** | **5.14** | **7.12** | **1** | **3** | **3** | **3** | **351** | **39.2** | **5.35** |
| B7QAH2 | Integrin α-ps, putative | 8038317 IscW_ISCW022321 | 327 | 4.98 | 9.48 | 1 | 2 | 2 | 2 | 327 | 35.5 | 5.27 |
| B7QBE1 | Guanylate cyclase (EC 4.6.1.2) | 8038704 IscW_ISCW012198 | 524 | 3.94 | 1.34 | 1 | 1 | 1 | 2 | 524 | 58.8 | 8.38 |
| B7QMY2 | High affinity nerve growth factor receptor, putative (EC 2.7.10.2) | 8043662 IscW_ISCW015459 | 707 | 3.75 | 4.53 | 1 | 3 | 3 | 3 | 707 | 77.7 | 6.83 |
| B7QKP6 | Type 3 inositol 1,4,5-trisphosphate receptor, putative | 8042698 IscW_ISCW023346 | 2569 | 2.27 | 1.36 | 1 | 2 | 2 | 2 | 2569 | 289.8 | 6.23 |
| B7PQT5 | Seven transmembrane receptor, putative | 8052318 IscW_ISCW018766 | 426 | 2.19 | 5.4 | 1 | 1 | 1 | 1 | 426 | 48.4 | 6.4 |
| B7PHS9 | Import receptor subunit tom20, putative | 8029163 IscW_ISCW004475 | 244 | 2.12 | 2.87 | 1 | 1 | 1 | 1 | 244 | 27.4 | 5.5 |
| B7PB29 | Excitatory amino acid transporter, putative | 8027235 IscW_ISCW017534 | 914 | 2.06 | 1.31 | 1 | 1 | 1 | 1 | 914 | 101.8 | 6.51 |
| B7Q593 | Serine/threonine-protein kinase receptor (EC 2.7.11.30) (Fragment) | 8036103 IscW_ISCW020983 | 452 | 2 | 2.21 | 1 | 1 | 1 | 1 | 452 | 50.7 | 6.4 |
| B7QH05 | FGF receptor activating protein, putative | 8041118 IscW_ISCW023531 | 263 | 1.94 | 3.04 | 1 | 1 | 1 | 1 | 263 | 29.8 | 8.88 |
| B7PGJ8 | Adenosine A3 receptor, putative (GPR insulin like receptor 2) | 8028650 IscW_ISCW018360 | 611 | 1.73 | 1.47 | 1 | 1 | 1 | 1 | 611 | 68.6 | 8.07 |
| B7PXC5 | Golgi SNAP receptor complex member 1 | 8032744 IscW_ISCW020319 | 240 | 0 | 3.75 | 1 | 1 | 1 | 1 | 240 | 27.3 | 7.34 |
| B7QAV0 | Receptor protein serine/threonine kinase (EC 2.7.11.30) (Fragment) | 8038463 IscW_ISCW022003 | 505 | 0 | 4.16 | 1 | 1 | 1 | 1 | 505 | 57.5 | 8.07 |

*Results were filtered using the Contaminant Repository for Affinity Purification (CRAPome) database to remove common contaminants. Table displays results subsequently filtered by the gene ontology (GO) term "membrane" and the keyword "receptor". PSMs=peptide spectrum matches, AAs=amino acids, MW=molecular weight, KDa=kilodaltons, pI=isoelectric point.*

Table S3. Constraint of *CD36* family members in the Genome Aggregation Database (gnomAD).

| Gene | CHR | POS | Expected LoF SNVs | Observed LoF SNVs | o/e (90% CI) |
|---|---|---|---|---|---|
| ***SCARB1*** | 12 | 125261402-125367214 | 25 | 7 | 0.028 (0.16-0.52) |
| ***SCARB2*** | 4 | 7079890-77135046 | 25 | 11 | 0.44 (0.27-0.72) |
| ***CD36*** | 7 | 79998891-80308593 | 23 | 66 | 2.8 (1.8-2) |

*SCARB1 and SCARB2 were highly constrained (o/e<1) when compared to CD36 (o/e>1). CHR=chromosome; POS=position in GRCh37/hg19; LoF=loss-of-function; SNVs=single nucleotide variants; expected LoF SNVs=expected number of LoF SNVs in gnomAD based on a mutational model that accounts for sequence context, coverage, and methylation; observed LoF SNVs=observed number of LoF SNVs in gnomAD; o/e (90% CI)=constraint score of the ratio of observed to expected number of LoF variants in the gene along with the 90% confidence interval.*

Table S4. Association of *CD36* loss-of-function variants with Lyme disease.

| Variant | CHR | POS | REF | ALT | Consequence | *rs*ID | N with variant (frequency) | OR | 95% CI | *P* Value |
|---|---|---|---|---|---|---|---|---|---|---|
| NC_000007.14:g.80664501_80664504dup | 7 | 80664497 | A | AAGTA | Splice donor | *rs*3211907 | 381 (0.013) | 3.4 | 1.3 - 7.6 | 0.019 |
| NC_000007.14:g.80656605C>G p.Y62Ter | 7 | 80656605 | C | G | Stop gained | *rs*574416705 | 5 (0.00017) | 73 | 7.1 - 417 | 0.0022 |
| NC_000007.14:g.80672839del/dup p.I399fs | 7 | 80672834 | A | *, AA | Frameshift | *rs*753754773 | 8 (0.00028) | 20 | 2.0 - 102 | 0.016 |

*Three CD36 LoF alleles with an allele frequency less than or approximately 1% were identified in BioMe and analyzed for Lyme disease diagnosis. Results showed the effect size of each variable on Lyme disease diagnosis. All three rare variants were significantly associated with Lyme diagnosis. Variant=variant identifier with reference sequence file identifier from NCBI/RefSeq; CHR=chromosome; POS=position in GRCh38/hg38; REF=non-effect allele; ALT=effect allele; Consequence=predicted molecular consequence using Variant Effect Predictor; rsID=variant identifier in Single Nucleotide Polymorphism Database (dbSNP); N with variant (frequency)=number of individuals with the variant and the variant frequency; OR=odds ratio adjusted for age, sex, body mass index, and 10 genetic principal components; 95% CI=95% confidence interval.*

Table S5. Association of *CD36* loss-of-function burden with total cholesterol.

| Variable | Estimate | SE | *P* Value |
|---|---|---|---|
| ***CD36* LoF burden** | **4.4** | **2.1** | **0.041** |
| Statin | -2.3 | 0.62 | $1.8 \times 10^{-4}$ |
| Age | -0.015 | 0.018 | 0.42 |
| Sex | -16 | 0.54 | $<2.0 \times 10^{-16}$ |
| BMI | 0.046 | 0.041 | 0.25 |

*Addition of a CD36 LoF allele was associated with a 4.4 mg/dL increase in total cholesterol. Variable=predictor variable in multivariable regression of cholesterol levels; CD36 LoF burden=burden of LoF variants in CD36 calculated as the sum of all CD36 LoF alleles present in a carrier (0=no LoF alleles, 1=one LoF alleles, 2=two LoF alleles); BMI=body mass index; estimate=effect size estimate of CD36 LoF burden on cholesterol levels adjusted for statin use, age, sex, body mass index, and 10 genetic principal components; SE=standard error.*

Table S6. Association of *CD36* loss-of-function (LoF) burden with Lyme disease.

| Variable | OR | 95% CI | *P* Value |
|---|---|---|---|
| ***CD36* LoF burden** | **2.1** | **1.3 - 3.1** | **$3.1 \times 10^{-4}$** |
| Age | 1 | 0.99 - 1.0 | 0.3 |
| Sex | 0.9 | 0.70 - 1.2 | 0.41 |
| BMI | 0.98 | 0.95 - 1.0 | 0.037 |

*Addition of a CD36 LoF allele is associated with a 2.1-fold increased odds for Lyme disease diagnosis. Variable=predictor variable in multivariable regression of Lyme disease; CD36 LoF burden=burden of LoF variants in CD36 calculated as the sum of all CD36 LoF alleles present in a carrier (0=no LoF alleles, 1=one LoF alleles, 2=two LoF alleles); BMI=body mass index; OR=odds ratio adjusted for age, sex, BMI, and 10 genetic principal components; 95% CI=95% confidence interval.*

Table S7. Association of *CD36* loss-of-function burden with Lyme disease accounting for cholesterol level.

| Variable | OR | 95% CI | *P* Value |
|---|---|---|---|
| ***CD36* LoF burden** | **2** | **1.3 - 3.0** | **$1.1x10^{-3}$** |
| **Cholesterol** | **1.004** | **1.0007 - 1.007** | **0.01** |
| Age | 0.99 | 0.98 - 1.0 | 0.071 |
| Sex | 0.8 | 0.60 - 1.1 | 0.12 |
| BMI | 0.97 | 0.94 - 0.99 | 0.0087 |

*Addition of a CD36 LoF allele is associated with a 2.0-fold increase odds for Lyme disease diagnosis when accounting for total cholesterol. Variable=predictor variable in multivariable regression of Lyme disease; CD36 LoF burden=burden of LoF variants in CD36 calculated as the sum of all CD36 LoF alleles present in a carrier (0=no LoF alleles, 1=one LoF alleles, 2=two LoF alleles); BMI=body mass index; OR=odds ratio adjusted for total cholesterol, age, sex, BMI, and 10 genetic principal components; 95% CI=95% confidence interval.*

Table S8. Baseline characteristics of study population from the Bio*Me* Biobank.

| Characteristic | Bio*Me* Biobank (*n*=28,877) |
|---|---|
| **Age, mean (SD)** | 58 (26) |
| **Sex, *n* (%)** | |
| **Female** | 16,539 (57) |
| **Male** | 12,338 (43) |
| **Ancestry, *n* (%)** | |
| **European** | 9,559 (33) |
| **African** | 6,847 (24) |
| **Hispanic** | 9,387 (33) |
| **Other** | 3,023 (10) |
| **Measurements, mean (SD)** | |
| **BMI, kg/m$^2$** | 28 (6.7) |
| **Total cholesterol, mg/dL** | 180 (39) |
| **Neutrophils, %** | 62 (11) |
| **Lymphocytes, %** | 27 (10) |
| **Statins, *n* (%)** | 8,368 (29) |
| **Diagnoses, *n* (%)** | |
| **CAD** | 6291 (22) |
| **Lyme disease** | 256 (0.89) |

*Ancestry, self-reported; other, other miscellaneous ancestries besides the ones listed; measurements, clinical and laboratory; BMI, body mass index; CAD, coronary artery disease. Values in parentheses denote standard deviation (SD) or percent (%) and are labeled accordingly.*

Table S9. Primers and siRNA sequences.

| Target | Type | Start Position Within mRNA | siRNA/Primer Name | Strand | Primer Sequence | Accession number |
|---|---|---|---|---|---|---|
| ***I. scapularis croquemort*** | siRNA (for *B. burgdorferi* acquisition)* | 513 | siCrq_513F | forward | AACCTTCCAGAGGGTCTATTTCCTGTCTC | XM_029984267.4; empirically determined sequence |
| | | | siCrq_513R | reverse | AAAAATAGACCCTCTGGAAGGCCTGTCTC | |
| | siRNA (for all other *in vitro* and *in vivo* experiments) | 1157 | siCrq_1157F | forward | AAGCATTCAAACCAACTTCATCCTGTCTC | |
| | | | siCrq_1157R | reverse | AAATGAAGTTGGTTTGAATGCCCTGTCTC | |
| | scrambled (for *B. burgdorferi* acquisition)* | | scCrq_513F | forward | AAGCTCGTGAACTCGTATTCTCCTGTCTC | |
| | | | scCrq_513R | reverse | AAAGAATACGAGTTCACGAGCCCTGTCTC | |
| | scrambled (for all other *in vitro* and *in vivo* experiments) | | scCrq_1157F | forward | AAGATCACAGAGTGCAAACTACCTGTCTC | |
| | | | scCrq_1157R | reverse | AATAGTTTGCACTCTGTGATCCCTGTCTC | |
| | qRT-PCR | 578 | Crq_F | forward | CCCAAGTTGCAGGAGGTGG | |
| | | 663 | Crq_R | reverse | CGCACCTCCCGGTAGG | |
| ***I. scapularis jun*** | siRNA (pooled) | 161 | siJun_161F | forward | AAGGATACGACATTCTACGAACCTGTCTC | XM_029975161.4 |
| | | | siJun_161R | reverse | AATTCGTAGAATGTCGTATCCCCTGTCTC | |
| | siRNA (pooled) | 782 | siJun_782F | forward | AAGGAGAGGATCAAGCTGGAACCTGTCTC | |
| | | | siJun_782R | reverse | AATTCCAGCTTGATCCTCTCCCCTGTCTC | |
| | scrambled (pooled) | | scJun_161F | forward | AAGATGCGACAAGATATCCTACCTGTCTC | |
| | | | scJun_161R | reverse | AATAGGATATCTTGTCGCATCCCTGTCTC | |
| | scrambled (pooled) | | scJun_782F | forward | AAGACGAAGCGGAGATTAGGACCTGTCTC | |
| | | | scJun_782R | reverse | AATCCTAATCTCCGCTTCGTCCCTGTCTC | |
| | qRT-PCR | 332 | Jun_F | forward | CGCAGTACCTGTTCACGAAG | |
| | | 497 | Jun_R | reverse | CGACGACGAGGGTAAGATGA | |
| ***I. scapularis relish*** | siRNA | 85 | siRelish_85F | forward | AACCTCACACCATTTGCCTTTCCTGTCTC | XM_040501061.2 |
| | | | siRelish_85R | reverse | AAAAAGGCAAATGGTGTGAGGCCTGTCTC | |
| | scrambled | | scRelish_85F | forward | AAGACGCCTATCGCCTACATTCCTGTCTC | |
| | | | scRelish_85R | reverse | AAAATGTAGGCGATAGGCGTCCCTGTCTC | |
| | qRT-PCR | 487 | Relish_F# | forward | CACGTGCACCGCCTCACCATGAAGG | |
| | | 595 | Relish_R# | reverse | AGAATGTCCGCCACCGTTTTTTTCTGC | |

| | | | | | | |
|---|---|---|---|---|---|---|
| ***I. scapularis dorsal*** | siRNA | 1249 | siDorsal_1249F | forward | AACCTCGAAATGATCCTCCTTCCTGTCTC | XM_029969643.3 |
| | | | siDorsal_1249 R | reverse | AAAAGGAGGATCATTTCGAGGCCTGTCTC | |
| | scrambled | | scDorsal_1249 F | forward | AAGCCCTTACTGACCTAATCTCCTGTCTC | |
| | | | scDorsal_1249 R | reverse | AAAGATTAGGTCAGTAAGGGCCCTGTCTC | |
| | qRT-PCR | 2184 | Dorsal_F | forward | CTACGTCTGCTGGTGTTGTG | |
| | | 2239 | Dorsal_R | reverse | ACCCAAGAATGACGGGCATA | |
| ***I. scapularis tak1*** | siRNA | 1591 | siTAK1_1591F | forward | AAGCGAGTCGATGCAGATCTTCCTGTCTC | XM_029994364.2 |
| | | | siTAK1_1591R | reverse | AAAAGATCTGCATCGACTCGCCCTGTCTC | |
| | scrambled | | scTAK1_1591F | forward | AAGACGTTACCCGGTGATGTACCTGTCTC | |
| | | | scTAK1_1591R | reverse | AATACATCACCGGGTAACGTCCCTGTCTC | |
| | qRT-PCR | 1978 | Tak1_F | forward | CGGCGAAAGAGTTCCATCTG | |
| | | 2121 | Tak1_R | reverse | GCGTTCAGAGTCGAACCAAA | |
| ***B. burgdorferi recA*** | qRT-PCR | 194 | RecA_F# | forward | GTGGATCTATTGTATTAGATGAGGCT | U23457.1 |
| | | 390 | RecA_R# | reverse | GCCAAAGTTCTGCAACATTAACACCT | |
| ***I. scapularis actin*** | qRT-PCR | 896 | I.s. actin_F† | forward | GGTCATCACAATCGGCAAC | XM_029977298 |
| | | 1003 | I.s. actin_R† | reverse | ATGGAGTTGTACGTGGTCTC | |
| ***I. scapularis puckered*** | qRT-PCR | 226 | Puckered_F | forward | GTCCTGTGGAGCTAAGGTGT | XM_040207225.2 |
| | | 307 | Puckered_R | reverse | TGGAAGACGTCGCCATAGAG | |
| ***I. scapularis ctenidin-1*** | qRT-PCR | 4 | Ctenidin1_F | forward | TTCCGTAGCAGCCACGTC | XM_029986701.1 |
| | | 60 | Ctenidin1_R | reverse | GCTGTGCCTGAATGACCAAG | |
| ***I. scapularis shade*** | qRT-PCR | 765 | Shade_F | forward | CAATGTCGGCCTCTCCAATG | XM_002401033.4 |
| | | 891 | Shade_R | reverse | AGCGCACATAGCTCACAAAG | |
| ***I. scapularis shadow*** | qRT-PCR | 4468 | Shadow_F | forward | AGACTTGTCACTGTCGCTCA | XM_002411796.4 |
| | | 4611 | Shadow_R | reverse | AAACACAAGCGGTCTACGTG | |
| ***I. scapularis disembodied*** | qRT-PCR | 1086 | Disembod_F | forward | AAGTTCGCACAGACATTCCG | XM_029973376.4 |
| | | 1234 | Disembod_R | reverse | TGGTGGCAAGGTAGTACAGG | |

| | | | | | | |
|---|---|---|---|---|---|---|
| ***I. scapularis ecr*** | qRT-PCR | 2840 | EcR_F | forward | CTCGCAGAGATCTGGGACAT | XM_040205666.3 |
| | | 3001 | EcR_R | reverse | TTGGCCGATTCGACTTTGAC | |
| ***I. scapularis usp*** | qRT-PCR | 1657 | USP_F | forward | GCCCTCAGGGAGAAGGTTTA | XM_002435025.5 |
| | | 1822 | USP_R | reverse | AGGAGGAAGTTGTCGATGGG | |
| **pET14b-Crq-ecto-His** | cloning | | pET14b-Crq-ecto-His_F | forward | GGCCCATATGATGAAGCTGTTCGTGAGCAT | XM_029984267.4 |
| | | | pET14b-Crq-ecto-His_R | reverse | GGCCCTCGAGGCTATAGTACAGCGGGCGGT | |
| **DsRed-N1-Crq** | cloning | | DsRed-N1-Crq_F | forward | CGTCAGATCCGCTAGCGGATGACATACTGCCGTGCTC | XM_029984267.4 |
| | | | DsRed-N1-Crq_R | reverse | CGTCGACTGCAGAATTCCCCCGTGAATAATGCCATTTTCCTTG | |

*This siRNA was designed prior to the existence of the newest *I. scapularis* genome.
[#]Shaw *et al.*, 2017; DOI:10.1038/ncomms14401
[†]Oliva Chavez *et al.*, 2021; DOI:10.1038/s41467-021-23900-8

Table S10. Resources and reagents available.

| Antibody | Source | Identifier | Dilution/Concentration |
|---|---|---|---|
| Goat Anti-Rabbit IgG (H+L) Cross-Adsorbed Secondary Antibody, Alexa Fluor 594 | Invitrogen | A-11012 | 1:1,000 |
| Rabbit (DA1E) mAb IgG XP® Isotype Control | Cell Signaling | 3900S | 0.111111111 |
| Rabbit Anti-*I.scapularis* Croquemort Polyclonal Ab | GenScript | custom | 1:100-1,000 |
| Rabbit Anti-Mouse Actin | Millipore Sigma | A2103 | 1:4,000 |
| Rabbit Phospho-SAPK/JNK (Thr183/Tyr185) Polyclonal Ab | Cell Signaling | 4668 | 1:1,000 |
| Rabbit JNK Polyclonal Ab | Proteintech | 10023-1-AP | 1:1,000 |
| Rabbit Anti-*I.scapularis* Relish Polyclonal Ab | GenScript | custom | 1:1,000 |
| $Na^+/K^+$-ATPase Rabbit Antibody | Cell Signaling | 3010S | 1:1,000 |
| GAPDH (14C10) Rabbit mAb | Cell Signaling | 2118S | 1:1,000 |
| Goat Anti-Rabbit IgG H&L (HRP) | Abcam | ab97051 | 1:4,000-10,000 |
| Mouse (G3A1) mAb IgG1 Isotype Control | Cell Signaling | 5415 | 5 mg |
| Mouse anti-*I.scapularis* N-Rel antiserum | GenScript | custom | 0.5-5 mg |
| Rat monoclonal [H139-52.1] Anti-Mouse kappa light chain (HRP) | Abcam | ab99632; discontinued | 1:4,000 |
| Phospho-NF-κB p65 (Ser536) (93H1) Rabbit mAb | Cell Signaling | 3033 | 1:1,000 |
| NF-κB p65 (D14E12) XP® Rabbit mAb | Cell Signaling | 8242 | 1:1,000 |
| **Cell Dye** | | | |
| CellMask™ Green plasma membrane stain | Invitrogen | C37608 | 1:1,000 |
| Hoechst 33342 Nuclear Stain | AAT Bioquest | 17533 | 1:1,000 |
| SlowFade Gold Antifade mountant | Invitrogen | S36938 | N/A |
| **Cell Media** | | | |
| Leibovitz's L-15 Medium, powder | Gibco | 41300039 | N/A |
| L-aspartic acid | Millipore-Sigma | 11189 | 0.449 g/L |
| L-glutamine | Millipore-Sigma | G8540 | 0.500 g/L |
| L-proline | Millipore-Sigma | 81709 | 0.450 g/L |
| L-glutamic acid | Millipore-Sigma | 49449 | 0.250 g/L |
| alpha-ketoglutaric acid | Millipore-Sigma | K1128 | 0.449 g/L |
| Sodium hydroxide | Millipore-Sigma | S8045 | 10 N |
| D-glucose | Millipore-Sigma | G7021 | 18.018 g/L |
| FBS (USDA approved; for tick media) | Millipore-Sigma | F0926-500ML | 10% |
| Bacto™ Tryptose Phophate Broth | BD | 260300 | 10% |
| Lipoprotein Concentrate | MP Biomedicals | 191476 | 0.1% |
| Normal rabbit serum | Pel-Freez | #31126-5 | 6.0% |
| Sodium bicarbonate | Millipore-Sigma | S6014 | 0.25% |
| HEPES | Millipore-Sigma | H4034 | 25 mM |
| CMRL1066 w/L-Glutamine (Powder) | US Biological | C5900 | N/A |
| Sodium citrate tribasic dihydrate | Millipore-Sigma | S4641 | 0.7 g/L |
| Yeastolate | BD | 255772 | 2 g/L |

| | | | |
|---|---|---|---|
| Neopeptone | BD | 211681 | 5 g/L |
| N-Acetyl-α-D-glucosamine | Millipore-Sigma | 1079-25GM | N/A |
| Albumin, Bovine Fraction V | MP Biomedicals | 160069 | N/A |
| Rifampicin | Millipore-Sigma | 557303 | 50 ug/ml |
| Phosphomycin | Millipore-Sigma | P5396 | 100 ug/ml |
| Amphotericin B | Gibco | 15290-026 | 1:100 |
| Sodium pyruvate | Millipore-Sigma | P5280 | 0.8 g/L |
| Distilled water | Gibco | 15-230-147 | N/A |
| LB broth | Millipore-Sigma | L3022 | N/A |
| LB agar with 100 ug/ml ampicillin | Quality Biological | 50-751-7582 | N/A |
| Ampicillin | Millipore-Sigma | A0166 | 100 ug/ml |
| Ammonium Chloride (15N, 99%) | Cambridge Isotope | NLM-467-10 | 0.6 g/L |
| MOPS minimal media | University of Maryland, Baltimore | N/A | N/A |
| DMEM, high glucose | Gibco | 11960044 | N/A |
| FBS (for BMDM culture) | Gemini Bio-products | 100-106 | 10% |
| Penicillin-Streptomycin Solution, 100x | Corning | 30-002-CI | 1X |
| **Materials** | | | |
| Cellstar® cell culture flasks, 25 $cm^2$ | Greiner bio-one | 690-160 | N/A |
| Cell culture plate with lid (6 well, flat bottom) | Millipore-Sigma | SIAL0516 | N/A |
| Costar® cell culture plate with lid (24 well, flat bottom) | Corning | CLS3526-1EA | N/A |
| NuPAGE™ 4-12% Bis-Tris Protein Gels, 1.5 mm, 10-well | Thermo Scientific | NP0335BOX | N/A |
| Mini-Protean® TGX™ gels | Biorad | 456-9034 | N/A |
| Trans-blot® Turbo™ Tranfer pack, 0.2 µm PVDF | Biorad | 1704156 | N/A |
| 5% Mini-PROTEAN® TBE Gel | Biorad | 4565014 | N/A |
| Biodyne B Precut Nylon Membranes | Thermo Scientific | 77016 | N/A |
| Amicon Ultra centrifugal filters (30K) | Millipore Sigma | UFC903024 | N/A |
| Miller GP 0.2 µm filter unit | Millipore-Sigma | SLGP033RS | N/A |
| FALCON® 14 ml Polypropylene round-bottom tube | Corning | 352059 | N/A |
| Nunc™ 96-Well Polystyrene Round Bottom Microwell Plates | Thermo Scientific | 262162 | N/A |
| 250 mm glass desiccator | Fisher Scientific | 08-615B | N/A |
| 1.5 ml microcentrifuge tubes | Thomas Scientific | 1148T71 | N/A |
| 15 ml conical screw cap tubes | USA Scientific | 5618-8261 | N/A |
| 50 ml conical screw cap tubes | USA Scientific | 5622-7270 | N/A |
| 500 ml vacuum filter/storage bottle system, 0.2 µm | Corning | 430773 | N/A |
| Rnase-free disposable pellet pestles | Fisher Scientific | 12-141-368 | N/A |
| 25 gauge, 5/8" needle | BD | 305122 | N/A |
| 27 gauge, 1/2" needle | BD | 305109 | N/A |
| Siliconized tips | VWR | 53503-800 | N/A |
| Nonstick microcentrifuge tubes | Ambion | AM12450 | N/A |
| Flowmi™ Cell Strainer | SP Bel-Art | 136800040 | N/A |
| Microscope slides | Fisher Scientific | 12550400 | N/A |

| | | | |
|---|---|---|---|
| **Reagents** | | | |
| Halt™ phosphatase inhibitor cocktail (100x) | Thermo Scientific | 78426 | 1:100 |
| Halt™ protease inhibitor cocktail (100x) | Thermo Scientific | 87786 | 1:100 |
| 10X RIPA | Millipore-Sigma | 20-188 | 1X |
| Chloroform | Millipore-Sigma | 288306 | 100% |
| Methanol anhydrous, 99.8% | Millipore-Sigma | 322415-1L | 100% |
| Ethyl alcohol, Pure; 200 proof for molecular biology | Millipore-Sigma | E7023-1L | 70-100% |
| Lipofectamine 3000 Reagent | Thermo Scientific | L3000008 | 7.5 ul/1 ml |
| TRIzol reagent | Ambion | 15596018 | N/A |
| iTaq™ Universal SYBR® Green Supermix | Biorad | 1725121 | 1X |
| Potassium nitrate | Fisher Scientific | P263-500 | N/A |
| Ketamine HCl | Henry Schein Animal Health | 56344 | 0.6 mg/20 g mouse |
| Xylazine | Akorn Animal Health | XYBAL | 0.06 mg/20 g mouse |
| Ethanol, 200 proof | Decon Labs | 2701 | 70% |
| POPG (16:0, 18:1 PG) | Avanti | 840457 | 10 ng/ml |
| Biotinylated POPG | Avanti | Custom | 50 ug |
| Streptavidin beads | Thermo Scientific | 88816 | 50 ul |
| 1X PBS | Quality Biological | 114-058-101 | N/A |
| HyClone™ Water, Molecular Biology Grade | Cytiva | SH3053801 | N/A |
| Isopropyl β-D-1-thiogalactopyranoside (IPTG) | Millipore-Sigma | I6758 | 0.5 mM |
| IGEPAL® CA-630 | Millipore-Sigma | I3021-100ML | 0.05% |
| Lysozyme from chicken egg white | Millipore-Sigma | L4919 | 0.2mg/ml |
| Guanadine HCl | Millipore-Sigma | G3272 | 6 M |
| Sodium Chloride | Millipore-Sigma | S7653 | 300 mM |
| Trizma® base | Millipore-Sigma | 93362 | 20 mM |
| Imidazole | Millipore-Sigma | I2399 | 150 mM |
| Hi60 Ni Superflow resin | Takara Bio | 635659 | N/A |
| Glycerol | Millipore-Sigma | G6279 | 10-50% |
| $D_2O$ | Cambridge Isotope | DLM-4-25 | 10% |
| 2-βmercaptoethanol | Gibco | 21985-023 | 5% |
| 6X Laemmli buffer | Alfa Aesar | J60660 | 1X |
| Blocking grade blocker, non fat skim milk | Biorad | 1706404 | 5% |
| Bovine Serum Albumin | Millipore-Sigma | A2058 | 3% |
| Sodium dodecyl sulfate (SDS) | Millipore-Sigma | L6026 | 0.10% |
| Paraformaldehyde | Millipore-Sigma | P6148 | 4.00% |
| GelCode™ Blue Stain Reagent | Thermo Scientific | 24590 | N/A |
| HBS-EP buffer | Activa | BR100188 | N/A |
| **Commercial Assays** | | | |
| Pierce™ Silver Stain for Mass Spectrometry | Thermo Scientific | 24600 | N/A |
| Pure Link RNA mini kit | Ambion | 12183025 | N/A |
| Silencer™ siRNA Construction Kit | Thermo Scientific | AM1620 | N/A |

| | | | |
|---|---|---|---|
| Verso cDNA Synthesis Kit | Thermo Scientific | AB-1453B | N/A |
| Pierce BCA Protein Assay Kit | Thermo Scientific | 23227 | N/A |
| Pierce ECL Western Blotting Substrate | Thermo Scientific | 32106 | N/A |
| DNeasy® Blood and Tissue kit | Qiagen | 69506 | N/A |
| QIAamp® DNA Mini Kit | Qiagen | 51304 | N/A |
| QIAprep® Spin Miniprep Kit | Qiagen | 27106 | N/A |
| *Mycoplasma* testing kit | Southern Biotech | 13100-01 | N/A |
| SF Cell Line 4D-NucleofectorTM X Kit L | Lonza Bioscience | V4XC-2012 | N/A |
| 20-Hydroxyecdysone EIA Kit | Cayman Chemicals | 501390 | N/A |
| NE-PER Nuclear and Cytoplasmic Extraction Reagents | Thermo Scientific | 78833 | N/A |
| LightShift™ Chemiluminescent EMSA Kit | Thermo Scientific | 20148 | N/A |
| IL-6 Mouse ELISA Kit | Invitrogen | KMC0061 | N/A |
| KC/CXCL1 Mouse ELISA Kit | Invitrogen | EMCXCL1 | N/A |
| Mouse IL-1 β/IL-1F2 Quantikine ELISA Kit | R&D Systems | MLB00C | N/A |
| **Equipment** | | | |
| Model 120 Sonic Dismembrator | Fisherbrand | FB120110 | N/A |
| Savant SpeedVac | Thermo Scientific | SVC100H | N/A |
| 4D-Nucleofector$^{TM}$ System | Lonza Bioscience | AAF-1002 | N/A |
| Nanoject III | Drummond Scientific Company | 3-000-207 | N/A |
| Cytospin 4 | Thermo Scientific | A78300003 | N/A |
| CFX96 Touch Real-Time PCR Detection System | Biorad | Discontinued | N/A |
| C1000 Touch Thermocycler | Biorad | 1851148 | N/A |
| **Cell Lines** | | | |
| *Ixodes scapularis* ISE6 cells | Ulrike Munderloh, University of Minnesota | ISE6 | N/A |
| *Ixodes scapularis* IDE12 cells | Ulrike Munderloh, University of Minnesota | IDE12 | N/A |
| **Organisms** | | | |
| *Ixodes scapularis* nymph ticks | Tick Lab, Oklahoma State University | N/A | N/A |
| *Ixodes scapularis* nymph ticks | Ulrike Munderloh, University of Minnesota | N/A | N/A |
| C57BL6J mice | University of Maryland, Baltimore | N/A | N/A |
| C57BL6J mice | Jackson Laboratories | #000664 | N/A |
| *Cd36*$^{-/-}$ mice | Jackson Laboratories | #019006 | N/A |
| C3H/HeJ mice | Jackson Laboratories | #000659 | N/A |
| *Escherichia coli* BL21(DE3) | Thermo Scientific | EC0114 | N/A |
| *Borrelia burgdoferi* B31 clone MSK5 | Jon Skare, Texas A&M University Health Science Center | N/A | N/A |

| | | |
|---|---|---|
| **Plasmids** | | |
| pET-14b | Novagen | 69660 |
| DsRed-N1 | Addgene | 54493 |
